# A Scube2-Shh feedback loop links morphogen release and spread to morphogen signaling to enable scale invariant patterning of the ventral neural tube

**DOI:** 10.1101/469239

**Authors:** Zachary M. Collins, Anna Cha, Albert Qin, Kana Ishimatsu, Tony Y.C. Tsai, Ian A. Swinburne, Pulin Li, Sean G. Megason

## Abstract

To enable robust patterning, morphogen systems should be resistant to variations in gene expression and tissue size. Here we explore how the Sonic Hedgehog (Shh) morphogen gradient in the ventral neural tube enables scaled patterning in embryos of varying sizes. Using zebrafish eggs that have been surgically reduced in size, we find that ventral neural tube patterning remains proportional in smaller embryos. Intriguingly, a secreted protein implicated in Shh release, Scube2, is expressed in the dorsal-intermediate neural tube far from Shh producing cells. Overexpression of *scube2* expands the Shh gradient whereas loss of *scube2* causes gradient contraction. Conversely, upregulation of Shh represses *scube2* expression while Shh downregulation increases *scube2* expression thus establishing a negative feedback loop. This regulatory feedback is necessary for scaling, as demonstrated by its loss in *scube2* overexpressing embryos. Using mathematical modeling, we show that feedback control on diffusion and release rates of Shh allows the morphogen gradient to be robust to differences in field length and Shh gene dosage. We conclude that Scube2 promotes release and diffusion of Shh to allow gradient scaling in an extension to the expander-repressor model.

**Summary Statement:** The Shh morphogen gradient can scale to different size tissues by feedback between Scube2 mediated release and diffusion of Shh and Shh based inhibition of Scube2 expression

## Introduction

When Lewis Wolpert first posed the “French Flag Problem”, he was seeking the answer to this fundamental question: What systems enable proportional patterning in embryos independent of embryo size? By the time Wolpert formalized this problem, developmental biologists had long known that embryos scale their patterning programs in response to changes in embryo size (Wolpert, 1969). For example, sea urchin larva pattern normally from a single isolated blastomere up to the four-cell stage and amphibian embryos can survive bisection and pattern proportionally at a reduced size (Driesch, 1892; Morgan, 1895; Spemann, 1938; Cooke, 1981). Significant scaling of pattern formation to tissue availability seems to be a near universal property of developing organisms. Yet, 50 years after Wolpert’s statement of the French Flag Problem, how morphogen gradients scale to pattern domains of varied sizes remains unclear in many systems.

Recent theoretical studies have proposed a variety of mechanisms that could account for scaling of morphogen-mediated patterning (Ben-Zvi and Barkai, 2010; Umulis and Othmer, 2013). Some ways of adjusting morphogen gradient to size involve tuning production, degradation, or distribution of the morphogen. For example, models show scaling occurs when morphogen flux is size-dependent or when a morphogen enhances its degradation (Umulis and Othmer, 2013). Scaling could also be achieved if concentrations at the source and sink are fixed (Čapek and Müller, 2019). There is support for this source-sink mechanism in zebrafish dorsoventral patterning (Zinski et al., 2017). Other proposed mechanisms involve size-dependent players that affect morphogen distribution: modulators that change degradation or diffusion of the morphogen with or without feedback from the morphogen (Ben-Zvi and Barkai, 2010; Umulis and Othmer, 2013; Shilo and Barkai, 2017; Čapek and Müller, 2019). One of these models involving feedback interaction is termed expander-repressor integral feedback control (Ben-Zvi and Barkai, 2010). In this model, a morphogen represses the expression of another secreted protein, known as the expander, that affects the range of the morphogen such as by increasing its diffusion rate or decreasing its degradation rate. In such models, morphogen signaling will expand until it has reached an encoded equilibrium. This equilibrium is controlled by the morphogen’s repression of the expander, thus enabling “measurement” of the size of the domain in need of patterning. The first biological example of this mechanism was proposed in Xenopus axial patterning. In this original model, Chordin expands BMP signaling by binding BMP ligand (ADMP) and allowing shuttling of BMP ligands toward the ventral side (Ben-Zvi et al., 2008). Subsequent experimental work implicated another factor, Sizzled, as playing a central role in scaling in a mathematically equivalent manner to the expander-repressor model (Ben-Zvi et al., 2014; Inomata et al., 2013). However, recent work argues against an expander-repressor mechanism for BMP gradient formation, and supports a source-sink mechanism focused on control of Chordin proteolysis with Sizzled being dispensable (Tuazon et al., 2020). Expander-like relationships have also been proposed to regulate scaling of Dpp gradients during wing disc growth (Ben-Zvi et al., 2011; Hamaratoglu et al., 2011), although several other mechanisms have more recently been reported (Averbukh et al., 2014; Zhu et al., 2020; Romanova-Michaelides et al., 2022). While expander-like feedback circuits have been suggested to lead to scale invariance from synthetic patterns in bacterial colonies to vertebrate axial patterning (Cao et al., 2016; Ben-Zvi et al., 2008; Inomata et al., 2013; Ben-Zvi et al., 2014), the biochemical details of how proposed expanders alter morphogen spread and to what extent the expander-repressor model fits experimental results compared with other models for scaling is still not clear in many systems.

Though scaling of early axis patterning following size reduction has been extensively studied, the molecular mechanisms through which tissues and organs subsequently scale their patterning have received less attention (Ben-Zvi et al., 2008; Inomata et al., 2013). Previously, scaling of patterning during organ growth has been considered in the fly wing disc, which grows remarkably in size while maintaining proportion (Averbukh et al., 2014; Ben-Zvi et al., 2011; Hamaratoglu et al., 2011). In vertebrates, the developing neural tube has been a powerful model to study morphogen-mediated patterning (Briscoe and Small, 2015). While neural tube patterning does not expand isometrically over time with growth, embryos of different species maintain consistent embryonic proportions in the face of significant variation in organ size during initial patterning (Kicheva et al., 2014; Uygur et al., 2016).

The vertebrate ventral neural tube is patterned by the morphogen Sonic Hedgehog (Shh; Marti et al., 1995; Roelink et al., 1995). Shh is produced by the notochord and floorplate and induces ventral cell fates over a long range in a dose-dependent manner (Briscoe et al., 2001; Zeng et al., 2001). Various modes of Shh transport have been reported including free diffusion, lipoprotein particles, extracellular vesicles, and along cytonemes (Petrov et al., 2017; Sanders et al., 2013). While mechanisms of Shh transport have long been disputed, biochemical evidence supports soluble Shh as a primary component of long-range signaling. However, Shh ligands are dually lipid-modified and are highly lipophilic (Pepinsky et al., 1998; Porter et al., 1996a; Porter et al., 1996b), and thus require mechanisms to be released from the membrane and move through extracellular space. Shh release was largely thought to be achieved by the protein Dispatched, but more recent work has identified Scube2 as an additional factor in promoting Shh release (Burke et al., 1999; Creanga et al., 2012; Kawakami et al., 2002; Tukachinsky et al., 2012). Furthermore, release of Shh from sender cells was shown to occur as a complex with Scube2, and coreceptors in receiving cells aid in the transfer of Shh from Scube2 to the receptor Patched1 in mammalian cell culture (Wierbowski et al., 2020).

Scube2 is a <u>S</u>ignal sequence containing protein with a <u>CUB</u> domain and <u>E</u>GF-like repeats. The role of Scube2 in Shh signaling was first identified from work using the zebrafish *you* mutant which corresponds to *scube2* (Hollway et al., 2006; Kawakami et al., 2005; van Eeden et al., 1996; Woods and Talbot, 2005; Yang et al., 2002). Interestingly, while *scube2* mutants have defects in ventral patterning, *scube2* is predominantly expressed in the dorsal and intermediate neural tube in both mice and zebrafish (Grimmond et al., 2001; Kawakami et al., 2005; Woods and Talbot, 2005). Additionally, epistasis experiments indicated that Scube2 acts upstream of Patched to stimulate Shh signaling (Woods and Talbot, 2005). This effect was also found to be cell-non-autonomous, as mosaic injection of *scube2* mRNA was capable of rescuing Shh-signaling defects over a long range (Hollway et al., 2006; Woods and Talbot, 2005). Studies in cell culture then demonstrated that Scube2 facilitates release of Shh from secreting cells cell non-autonomously (Creanga et al., 2012; Tukachinsky et al., 2012). Recent work concluded that Scube2 may be responsible for catalyzing the shedding of lipids from Shh ligands, but this model is disputed by findings that released Shh remains dually lipid-modified and that a Scube2-Shh complex was shown to interact via lipid modifications on Shh (Creanga et al., 2012; Jakobs et al., 2014; Jakobs et al., 2016; Tukachinsky et al., 2012; Wierbowski et al., 2020). Biochemical evidence for the cell non-autonomous role of Scube2 in Shh release and its unexpected expression pattern in the zebrafish neural tube led us to wonder whether Scube2 may regulate pattern scaling by acting as an expander, as also hypothesized elsewhere (Shilo and Barkai, 2017) or in an expander-like role. In this work, we use quantitative imaging, genetic perturbations, and mathematical modeling to investigate the scaling of ventral neural patterning in size-reduced zebrafish embryos. We propose that Scube2 scales the Shh morphogen gradient with tissue size through feedback regulation of both morphogen release and diffusion in an extension to the expander-repressor model.

## Results

### Ventral neural patterning scales with embryo size

Studying mechanisms of scaling during growth or between species of different sizes is difficult because many properties of the patterning system depend on stage or species-specific variables. To study scaling of pattern formation in embryos with comparable genetic backgrounds at matched time points, we developed a technique to reduce the size of zebrafish embryos inspired by classical work in amphibians, as we previously described (Ishimatsu et al., 2018, 2019; Morgan, 1895; Spemann, 1938). Two lateral cuts are made across the blastula stage embryo: one to remove cells near the animal pole, to avoid damaging signaling centers crucial to early D-V patterning, and a second to remove yolk near the vegetal pole. (Fig. 1A). With this technique, a significant fraction of embryos patterns normally and develop at a reduced size (Fig. 1B-C).

**Figure 1.**
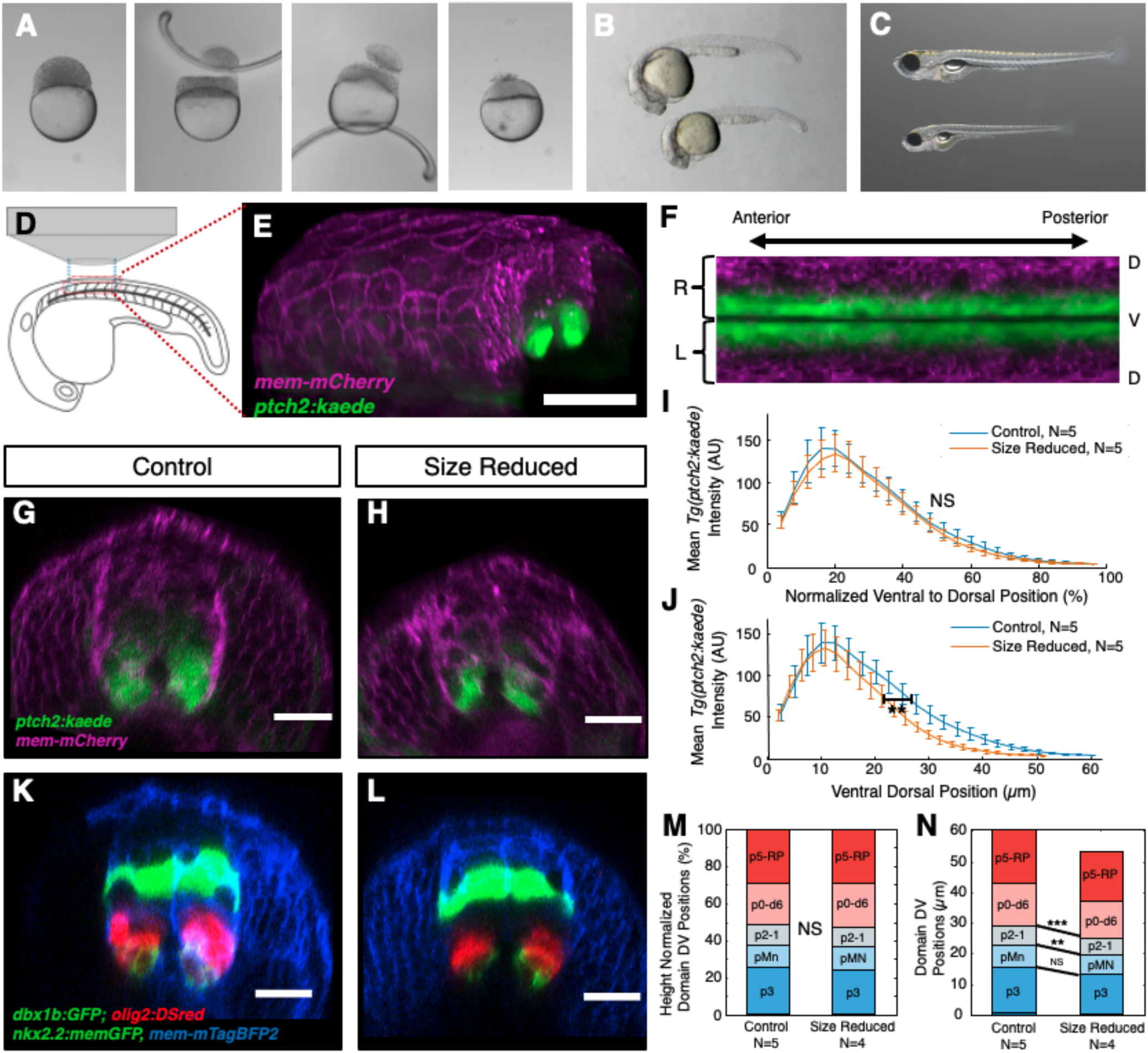
Neural tube patterning scales following embryonic size reduction. (A) Surgical size reduction of 128-256 cell stage embryos in which cells and yolk are removed to produce smaller embryos (adapted from Ishimatsu et al. 2018). (B) Size-reduced embryo at 24 hpf (lower) with a normal-sized sibling (upper). (C) Size-reduced larva at six days post fertilization (lower) with a normal-sized sibling (upper). (D) Schematic of an embryo mounted for imaging; anterior-posterior length of the imaging window is shown with blue lines. Red lines indicate the 3-D extent of the imaging window. (E) 3-D rendering of a confocal z-stack of *Tg(ptch2:kaede) mem-mCherry* mRNA injected embryo. Scale bar represents 100 μm. (F) Flattened “fillet” profile of segmented imaging data from a z-stack imaged as in E (See methods). (G,H,K,L) Scale bar represents 20 μm. (G-H) Transverse view of 20 hpf *Tg(ptch2:kaede*), *mem-mCherry* mRNA injected control (G) and size-reduced embryos (H). Size reduction led to a decrease in DV height of 15.0% +/- 2.8% in this dataset. (I-J) *ptch2:kaede* intensity profiles from segmented imaging data of embryos from G-H plotted either normalizing for D-V neural tube height (I) or to their absolute neural tube heights (J). Error bars indicate standard deviation. No significant shift is observed in the position of 50% mean maximum control intensity on a relative scale between control and size-reduced embryos(I) (unpaired t-test p=0.5981) while a statistically significant shift in the position is observed when comparing absolute positional values (J) (unpaired t-test p=0.0076; Control N=5, Size Reduced N=5). (K-L) Transverse views of *mem-mtagBFP2* mRNA injected 24 hpf *Tg(nkx2:mGFP; olig2:dsred; dbx1b:GFP*) control (K) and size-reduced (L) embryos. Size reduced embryos had an average 12.2% +/- 2.4% reduction in neural tube DV height relative to control embryos in this dataset. (M-N) Progenitor domain segmentation analysis (See methods) of embryos treated as in K-L plotted either normalizing for differences in neural tube height (relative scale) (M) or with respect to their absolute heights (absolute scale) (N). Only when compared on an absolute scale are statistically significant shifts seen in the dorsal boundaries of the p2-1 and pMN domains (absolute heights unpaired t-test p^p2-1^=1.0172e-4 and p^pMN^ =0.0016, Control N=5, Size Reduced N=4). Thresholds for P value significance asterisks: significant with p < 0.0001 noted with ****, with p = 0.0001 to 0.001 noted with ***; with p = 0.001 to 0.01 noted with **; with p = 0.01 to 0.05 noted with *.

We measured scaling of neural patterning in size-reduced embryos using quantitative imaging (Megason, 2009; Xiong et al., 2013). High-resolution image-stacks of 18-24 hours postfertilization (hpf) stage-matched zebrafish embryos were collected under identical settings, during the same imaging session, from matched Anterior-Posterior positions in control and experimentally perturbed embryos (Fig. 1D-E). Imaging volumes were analyzed with custom software to demarcate the dorsoventral axis and width of the neural tube along the length of the dataset (Fig. 1F, Fig. S1). Image intensity values were extracted in a set number of bins along the D-V axis for the left and right halves of the neural tube to normalize for variability in neural tube height. This system allowed for the quantitative and unbiased comparison of 3-4 somite lengths of neural imaging data from multiple embryos.

We quantified the Shh response gradient based on the expression of *patched2,* a direct transcriptional target of Shh, using the *Tg(ptch2:kaede*) reporter in wild type and size-reduced embryos (Fig. 1G-J) (Huang et al., 2012). When quantified relative to their respective neural tube dorsal-ventral heights, *Tg(ptch2:kaede*) response gradients maintained nearly identical intensity distributions despite the neural tube height being 15.0% (+/- 2.8%) smaller in size-reduced embryos in this dataset (N=5), indicating that Shh responses scale following size reduction (Fig. 1I-J). When viewed on an absolute scale, control and size-reduced embryos show clear shifts in the response gradient as measured by the position at which 50% of mean control maximum intensity is reached (p=0.0076) (Fig. 1J). To quantify this effect at the level of cell fate specification, we utilized a triple transgenic reporter line to simultaneously mark three major ventral neural fates: *nkx2.2a* (p3 progenitors), *olig2* (pMN and some p3 progenitors), and *dbx1b* (p0, d6 progenitors) (Fig. 1K-L) (Gribble et al., 2007; Jessen et al., 1998; Kinkhabwala et al., 2011; Kucenas et al., 2008). Anterior-posterior averaged intensity profiles were then segmented to form cell fate profiles which can be compared between embryos (Fig. S2). After normalizing for their altered D-V height (which was reduced in this group by 12.2% +/- 2.4% compared to controls), the average of these cell fate profiles in size-reduced embryos were virtually indistinguishable from those of full-sized embryos (Fig. 1M). Furthermore, differences between progenitor domain boundary positions were visible when size normalization was removed (Fig. 1N). Statistically significant shifts in the positions of the p2 and pMN upper boundaries were observed only when compared in their absolute coordinates (Fig. 1N). This further demonstrates that ventral neural patterning adjusts to changes in total D-V height.

### Scube2 levels control Shh signaling

Based on its role in the cell-non-autonomous regulation of Shh release and its dorsal expression pattern, we hypothesized a potential role for Scube2 in enabling scaling of Shh gradients (Kawakami et al., 2005; Woods and Talbot, 2005). This hypothesis depends on *scube2* expression levels having a dose-dependent effect on Shh signaling. However, previous work concluded that Scube2 is only required for Shh signaling as a permissive factor (Kawakami et al., 2005; Woods and Talbot, 2005). To examine the role of Scube2 in ventral neural patterning, we performed a morpholino knockdown of *scube2* in *Tg(ptch2:kaede*) reporter embryos using a previously validated translation inhibiting morpholino (Fig. 2A-C) (Woods and Talbot, 2005). We observed a decrease in Shh signaling following morpholino injection, as demonstrated by a statistically significant suppression of maximum *ptch2:kaede* intensity (Fig. 2C). Such a decrease in the amplitude of Shh signaling is consistent with a role of Scube2 in Shh release. Additionally, quantification of *nkx2.2a, olig2*, and *dbx1b* domain sizes in embryos injected with *scube2* morpholino showed a contraction of ventral progenitor domains (Fig. 2D-F). Ventral shifts in the upper boundaries of the pMN and p3 domains were statistically significant, due in part to near complete elimination of the *nkx2.2a+* p3 domain (Fig. 2F). Unexpectedly, expansion of the p2-1 domain dorsally was also observed, potentially implying long range Shh signaling is required for complete p2-1 induction.

**Figure 2.**
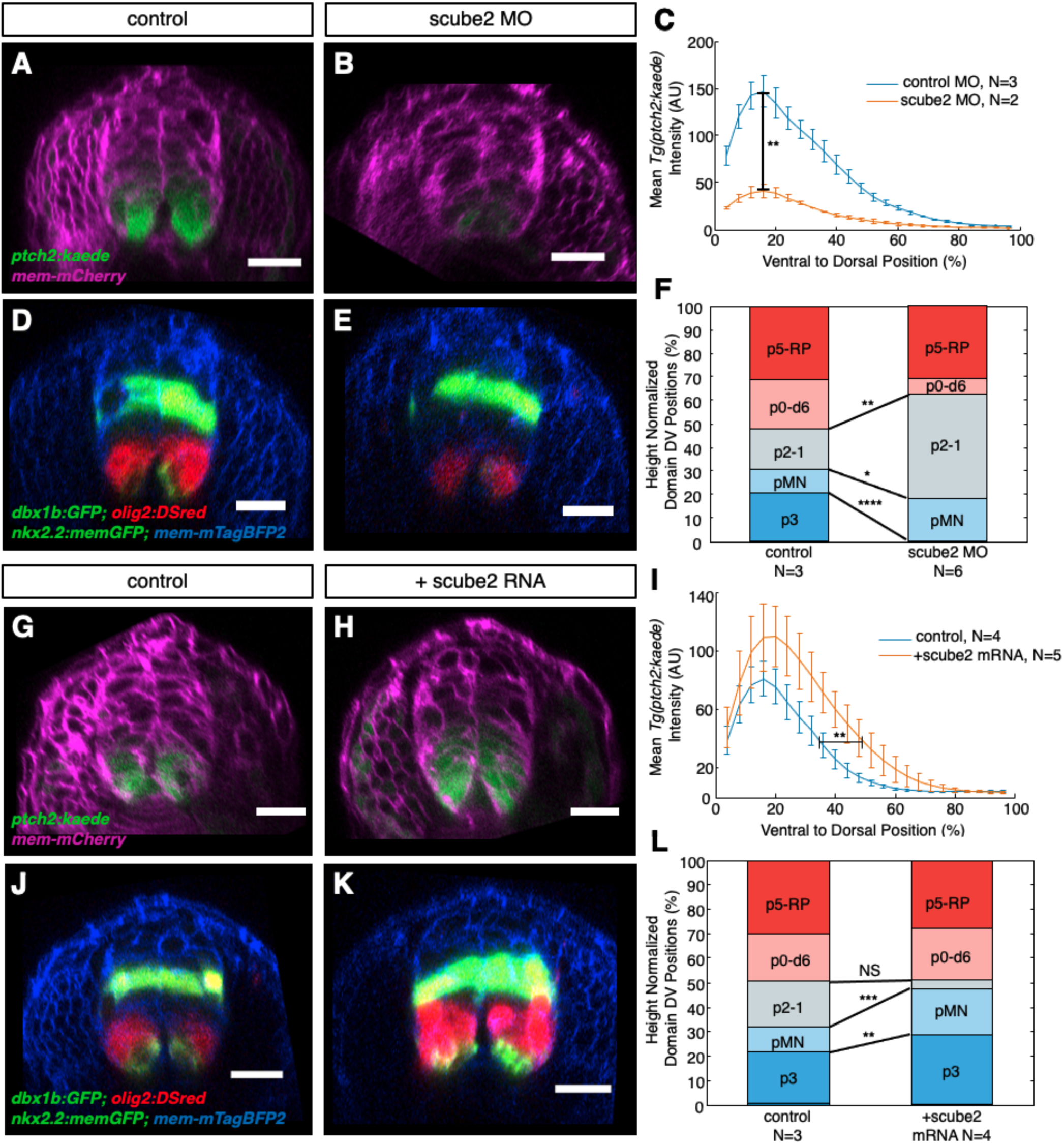
Scube2 expression levels regulate Shh signaling in the ventral neural tube. (A-B,D-E, G-H, J-K) Transverse view of a confocal z-stack, scale bar represents 20 μm (A-B) 22 hpf *Tg(ptch2:kaede*) embryos injected with (A) *mem-mCherry* mRNA with a control morpholino or (B) injected with *mem-mCherry* mRNA and *scube2* morpholino. (C) Mean intensity distributions in segmented neural tissue from z-stacks of embryos as treated in A-B. Maximum intensities of morpholino treated embryos were statistically significantly reduced compared to controls (p= 0.0040). (D-E) 20 hpf *Tg(dbx1b:GFP, olig2:dsred, nkx2.2a:memGFP*) embryos injected with (D) *mem-mTagBFP2* mRNA alone or (E) co-injected with *scube2* morpholino. (F) Mean result of automated segmentation of progenitor domain sizes (see methods for details) for embryos treated as in D-E. p1-2 domain upper boundaries were shifted dorsally following morpholino treatment (p^p2-1^= 0.0019), while the upper boundary of pMN and p3 domains were both significantly contracted in morpholino injected embryos (p^pMN^= 0.0158 and p^p3^= 9.87e-9). (G-H) Representative image of 20 hpf *Tg(ptch2:kaede*) reporter line embryos injected with (G) *mem-mcardinal* mRNA alone or (H) co-injected with *scube2* mRNA. (I) Quantification of mean intensity distributions of embryos as treated in G-H. Dorsoventral position at which half of average control maximum intensity was reached was significantly shifted in Scube2 overexpressing embryos (unpaired t-test p=0.00470). (J-K) 20 hpf *Tg(dbx1b:GFP, olig2:dsred, nkx2.2:mGFP*) embryos injected with (J) *mem-mTagBFP2* mRNA alone or (K) co-injected with *scube2* mRNA. (L) Mean results of automated progenitor domain segmentation of J-K. pMN and p3 domains were drastically shifted dorsally in *scube2* mRNA injected embryos (p^MN^=8.6748e-04, and p^p3^=0.0034 respectively).

Previous work concluded that Scube2 was a permissive factor based on a lack of change in expression of downstream Shh signaling markers as observed by whole mount *in situ* hybridization following *scube2* overexpression (Woods and Talbot, 2005). However, quantitative imaging reveals that injection of *scube2* mRNA leads to the expansion of Shh signaling, as shown by broader distributions of *Tg(ptch2:kaede*) fluorescence (Fig. 2G-I). Embryos injected with *scube2* mRNA showed significant dorsal shifts in the position of half maximum *Tg(ptch2:kaede*) intensity of the control (p=0.0047), a measure of absolute Shh signaling range (Fig. 2H). Maximum *Tg(ptch2:kaede*) fluorescence in *scube2* overexpressing embryos appeared higher on average but was not statistically significant (p=0.0512). In addition, *scube2* overexpression affected cell type patterning in the ventral neural tube, as measured in triple transgenic *nkx2.2a, olig2,* and *dbx1b* reporter embryos (Fig. 2J-L). Quantification of these cell fate profiles revealed large increases in p3 and pMN domain sizes, a decrease in the size of the p2-1 domains, and unchanged patterning of the p0-d6 domains and more dorsal cell types. Ventralization was measured by comparing the dorsal boundaries of the p3 and pMN, which were statistically significantly shifted (Fig. 2L). These data indicate that not only is Scube2 required for long range Shh signaling, but that *scube2* overexpression amplifies endogenous Shh signaling. Additionally, this suggests that Scube2-stimulated Shh release is a limiting factor in normal patterning.

### Shh negatively regulates Scube2 expression over a long-range

To study the regulation of Scube2 expression, we developed the *Tg(scube2:moxNG*) reporter line containing 7.6KB of the endogenous regulatory sequences driving moxNeonGreen fluorescent protein (Fig. 3A) (Costantini et al., 2015). The expression of *Tg*(*scube2:moxNG*) we observed is consistent with previously reported *in situ* hybridizations (Grimmond et al., 2001; Kawakami et al., 2005; Woods and Talbot, 2005). *Tg(scube2:moxNG*) embryos showed very low expression close to the sources of Shh in the floor plate and notochord—as visualized with a transgenic *Tg(shha:memCherry)* reporter line—and high levels of expression in the dorsal-intermediate neural tube (Fig. 3A-C). Time lapse imaging of *Tg(scube2:moxNG*) embryos revealed weak mesodermal expression in the early embryo, which faded during the onset of neurulation and was replaced by high levels of expression in the dorsal and intermediate neural tube (Fig. 3D-G, Movie S1-2).

**Figure 3.**
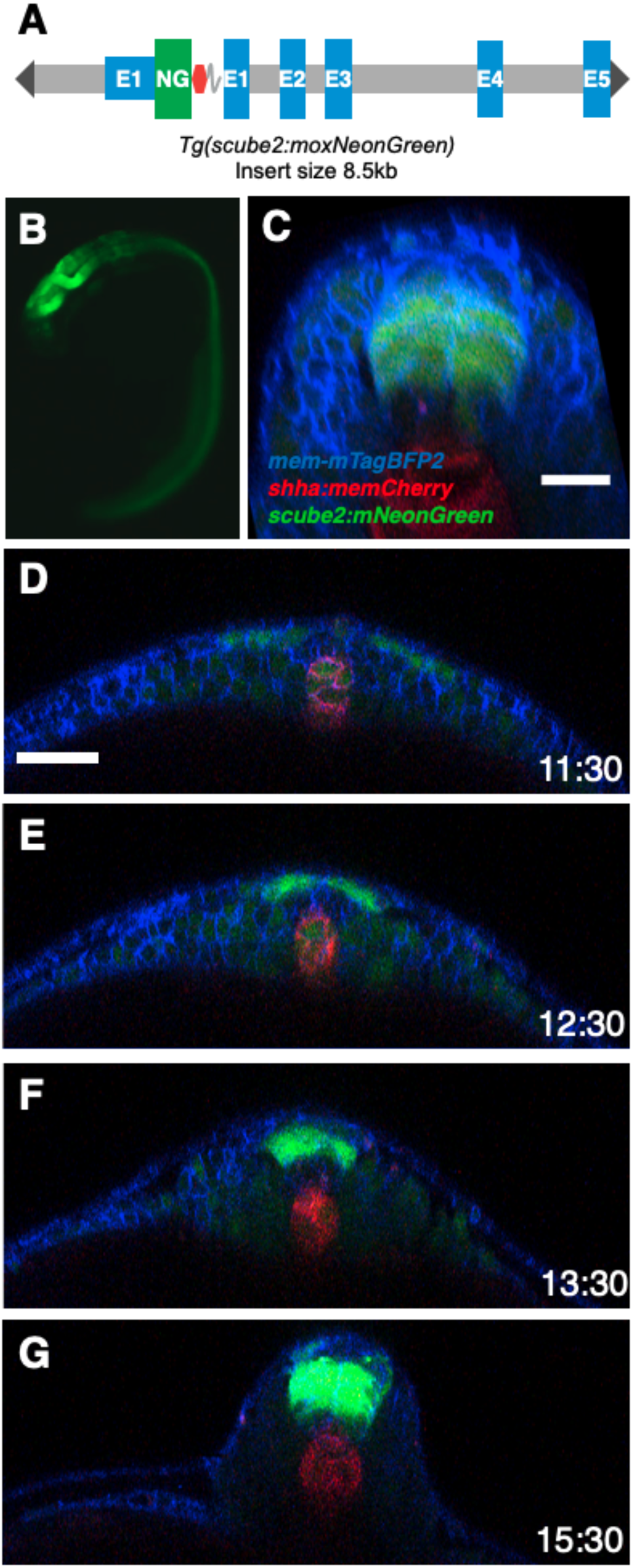
Scube2 is expressed distantly from Shh secreting cells. (A) Schematic of the *scube2:moxNG* transgenic expression reporter construct used to generate the *Tg(scube2:moxNG)* line. (B) Wide-field fluorescence image of *Tg(scube2:moxNG)* embryo at 20 hpf. (C) Transverse view of *mem-mTagBFP2* mRNA-injected *Tg(scube2:moxNG; shha:mem-mCherry)* embryo at 20 hpf. Scale bar represents 20 μm. (D-G) Transverse view from a time-lapse imaging dataset of *Tg(scube2:moxNG; shha:memCherry)* embryo which was injected at the single cell stage with *mem-mTagBFP2* mRNA. Time in hours post fertilization is displayed in the bottom right corner. Scale bar represents 50 μm. (D) At early neurulation stages there is weak mesodermal expression of *scube2:moxNG* in the notochord. In addition, expression of *scube2:moxNG* is visible in neural progenitors as the neural plate converges. (E) By 12.5 hpf a pronounced gap in expression of neural progenitors between *shha:mem-mCherry* and *scube2:moxNG* cells is visible. (F) Expansion of the *scube2*+ domain dorsally is visible as cells continue to converge. (G) *scube2* expression is constricted to the dorsal-intermediate neural tube.

To test whether Shh signaling downregulates Scube2 expression, we injected mRNA encoding a potent activator of Shh signaling, dnPKA, at the single cell stage (Hammerschmidt et al., 1996). Embryos injected with *dnpka* mRNA showed near complete ablation of neural *Tg(scube2:moxNG*) expression (Fig. 4 A-C). To test whether Shha ligands themselves were capable of suppressing Scube2 expression at a distance, we mosaically overexpressed *shha* in *Tg(scube2:moxNG*) embryos by injecting a single blastomere at the 16-cell stage with either *mem-mTagBFP2* alone or with *shha* mRNA (Fig. 4 D-F). We expected local inhibition of Scube2 reporter activity near secreting cells within a few cell diameters. Surprisingly, *Tg(scube2:moxNG*) expression was nearly completely eliminated in these embryos indicating potent cell-non-autonomous repression of Scube2 by Shh. When quantified, these embryos demonstrate a significant reduction of peak *Tg(scube2:moxNG)* intensities (Fig. 4F). To test whether Shh’s inhibition of Scube2 is required for its endogenous low ventral expression, we treated embryos with sonidegib, a potent Smoothened antagonist starting at the dome stage. Resulting embryos showed expanded *scube2* expression towards the floor plate and notochord (Fig. 4G-I). Shifts in ventral boundaries were statistically significant, as quantified by the D-V position at which half maximum intensity of the control population was reached (Fig. 4I). A ventral expansion *Tg(scube2:moxNG)* was also observed following cyclopamine treatment, which is consistent with the previous findings of a genome-wide screen for genes regulated by Shh signaling (Fig. S3) (Xu et al., 2006). These results indicate that endogenous Shh signaling is responsible for a lack of ventral *scube2* expression.

**Figure 4.**
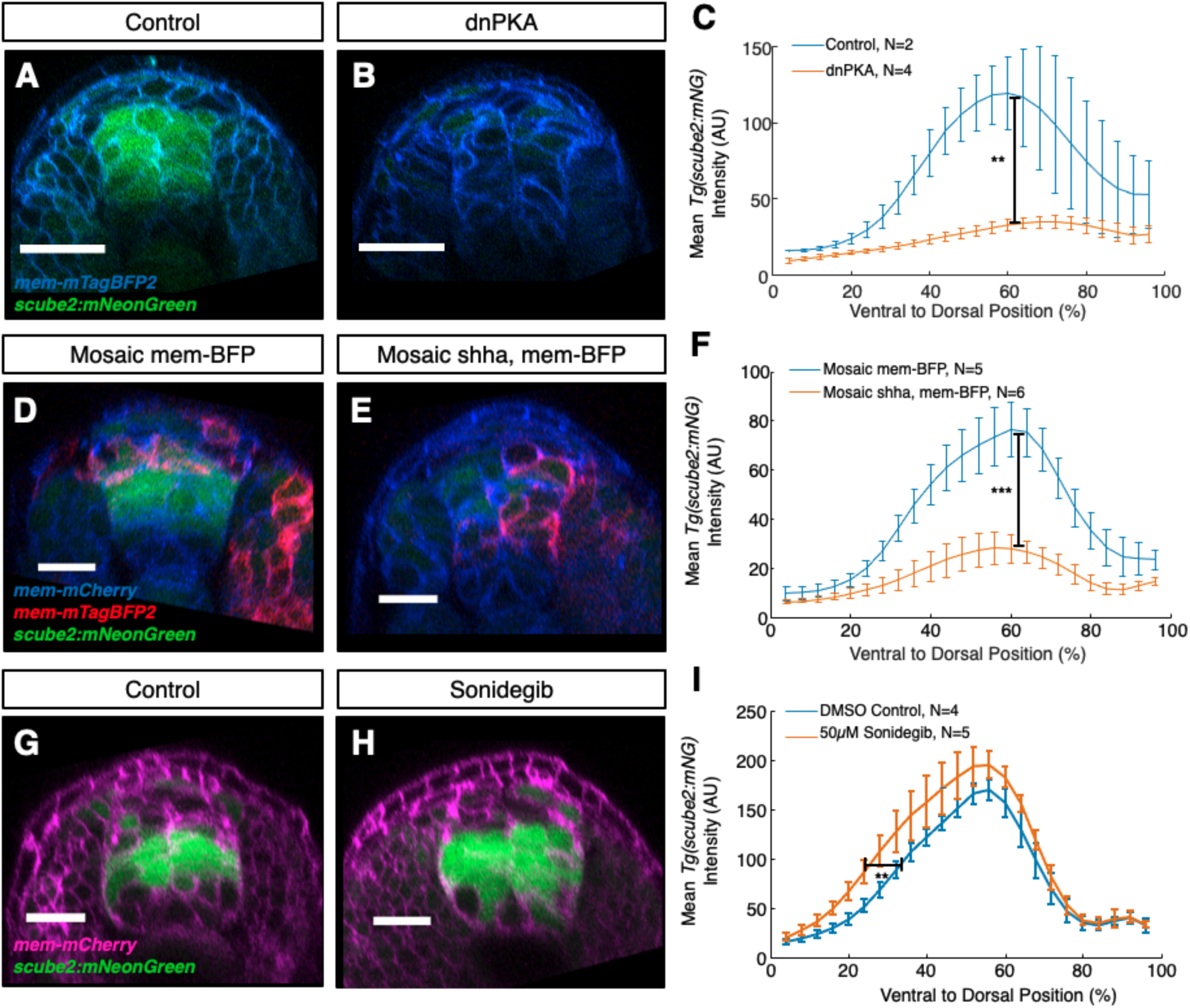
Shh signaling represses Scube2 expression. (A-B, D-E, G-H) Scale bar represents 20 μm (A-B) Transverse view of 18 hpf *Tg(scube2:moxNG)* embryos injected with (A) *mem-mTagBFP2* mRNA alone or (B) co-injected with *dnpka* mRNA. (C) Quantification of mean reporter intensity of embryos as treated in A-B. Maximum *scube2:moxNG* intensity values were significantly reduced in *dnpka* mRNA*-*injected embryos (p= 0.0014). (D-E) Transverse view 20 hpf *Tg(scube2:moxNG)* embryos injected at the single cell stage with *mem-mCherry* mRNA and then injected in one blastomere at the 8-16 cell stage with either (D) *mem-mTagBFP2* mRNA alone or (E) co-injected with *shha* mRNA. (F) Quantification of mean reporter intensity of embryos as treated in D-E. Maximum *scube2:moxNG* reporter intensity is significantly reduced in *shha* injected embryos (p= 9.14e-06). (G-H) Transverse view 22 hpf *Tg(scube2:moxNG*; *mem:mCherry)* embryos treated with a DMSO control (G) or treated with 50 μm sonidegib (H). (I) Quantification of mean reporter intensity of embryos as treated in G-H. The black bracket marks the position of half control maximum intensity used for statistical testing. These values were significantly shifted ventrally in drug treated embryos relative to control (unpaired t-test p=0.0014.).

To further probe the transcriptional regulation of Scube2’s expression we performed a small scale CRISPR mutagenesis screen and found that Pax6a/b are necessary for driving Scube2 expression. Co-injection of *pax6a* and *pax6b* sgRNAs with Cas9 caused significant downregulation of *Tg(scube2:moxNG)* relative to control embryos injected with a sgRNA targeting *tyrosinase,* an unrelated pigment gene (Fig. S4). These data support a model in which the expression domain of *scube2* is based on activation by Pax6a/b and repression by Shh.

### Scube2 diffuses over long distances during patterning

While Scube2 is known to act cell non-autonomously from transplantation experiments and Scube2-conditioned media has a potent Shh release stimulating effect *in vitro*, the localization of Scube2 protein during development is unknown (Woods and Talbot, 2005; Creanga et al., 2012). *In vitro*, Scube2 is thought to associate with Heparan Sulfate Proteoglycans, and Scube2 had been hypothesized to diffuse from secreting cells in the dorsal neural tube, the need for which was later disputed (Jakobs et al., 2016; Kawakami et al., 2005; Hollway et al., 2006). To examine Scube2’s localization, we developed Scube2 fluorescent fusion proteins by tagging the C-terminus as previously validated in cell culture with other tags (Fig. 5A) (Creanga et al., 2012). The resulting Scube2-mCitrine fusion proteins were functional and rescued Scube2 CRISPR mutants at comparable rates to wildtype Scube2 (Fig. S5). Mosaic injection of *scube2-mCitrine* mRNA into 1 cell at the 32-64 cell stage revealed that Scube2-mCitrine diffuses distantly from producing cells (Fig. 5B) and does not remain associated with cell membranes, as demonstrated by its presence in the extracellular space between cells marked with mem-mCherry (Fig. 5C).

**Figure 5.**
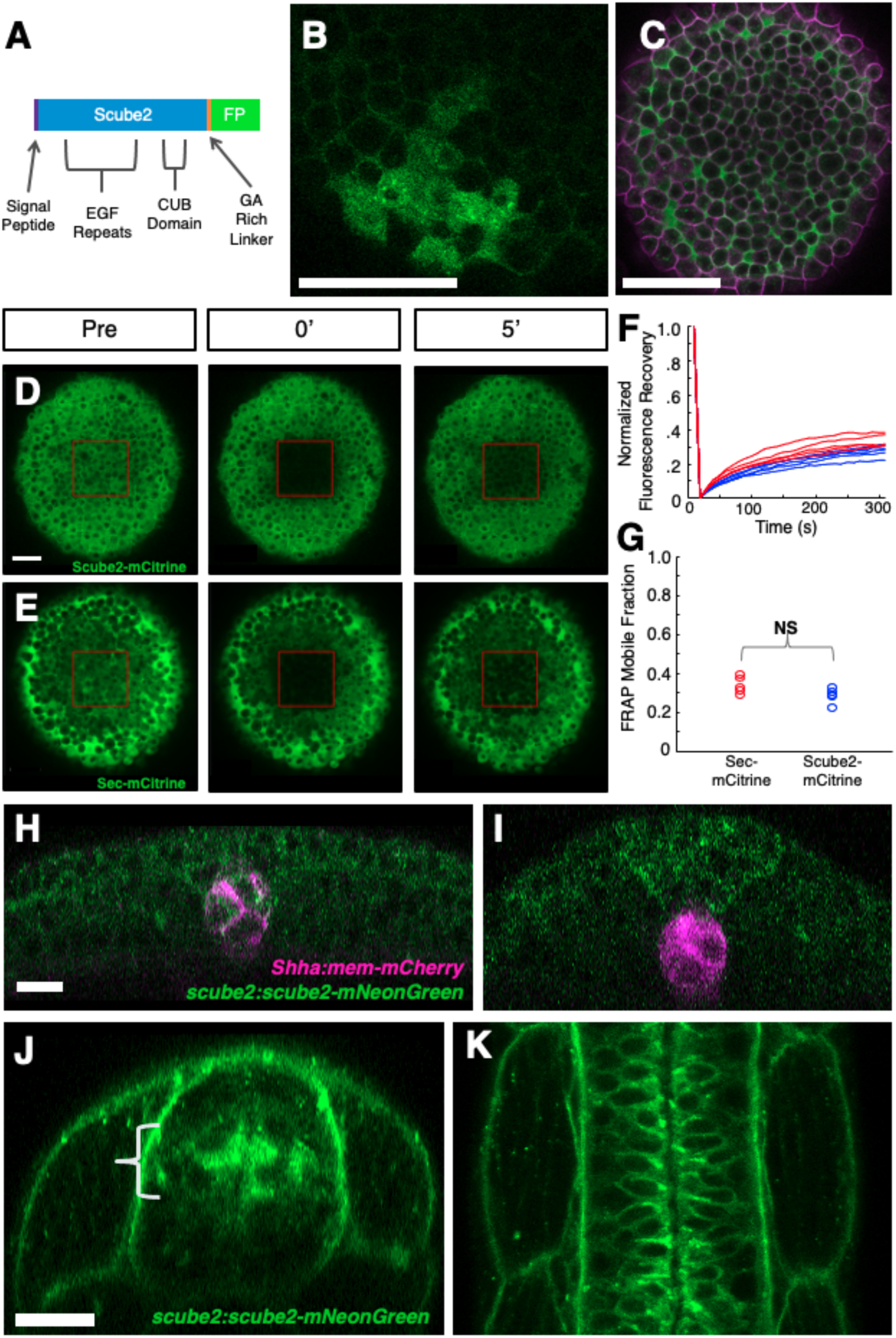
Scube2 diffuses from secreting cells and is broadly distributed during patterning. (A) Schematic of Scube2-mCitrine fluorescent fusion protein design. (B-C) Scale bar represents 100 μm. (B) Scube2-mCitrine fluorescence at the sphere stage from embryos injected in one blastomere at the 64-cell stage with *scube2-mCitrine* mRNA. Cells from the injected clone are marked by intracellular (secretory system) fluorescence. (C) Scube2-mCitrine fluorescence from embryos injected at the single cell stage with *scube2-mCitrine* and *membrane-mCherry* mRNA. (D-E) Fluorescence recovery after photobleaching at the dome stage of Scube2-mCitrine (D) and Secreted-mCitrine (E). (F) FRAP recovery traces normalized to maximum intensity pre-bleach and minimum intensity following bleaching. Red lines represent Secreted-mCitrine while blue lines represent Scube2-mCitrine. (G) Comparison of calculated mobile fraction values from dome stage FRAP data (unpaired t-test p=0.0698). (H-K) Scale bar represents 20 μm. (H) Transverse view of an 11.5 hpf *Tg(scube2:scube2-moxNG;shha:mem-mCherry)* embryo. (I) Transverse view of a 14 hpf *Tg(scube2:scube2-moxNG; shha:mem-mCherry)* embryo. (G) Transverse view of a 24 hpf *Tg(scube2:scube2-moxNG)* embryo. (K) Horizontal view of the embryo from J.

To assay Scube2’s rate of diffusion we performed Fluorescence Recovery After Photobleaching (FRAP) at the dome stage, during which cell movement is minimal. FRAP was performed in a 100µm x 100µm region and recovery was observed at 10 second intervals over 5 minutes (Fig. 5 D-E, Movie S3-4). Scube2-mCitrine fluorescence recovers almost as quickly as a secreted version of mCitrine generated using the Scube2 signal sequence (Fig. 5 F-G, see methods). These data indicate that Scube2 diffuses rapidly in the extracellular space, at a comparable rate to secreted mCitrine alone.

To observe distributions of Scube2 protein during development, we generated a transgenic line expressing the full-length Scube2 protein fused to moxNeonGreen under control of Scube2 regulatory sequences (Fig. 5H-K). We validated the functionality of this *Tg(scube2:scube2-moxNG)* line using a morpholino which bound only endogenous *scube2* RNA at the splice junction of exon6, and not *Tg(scube2:scube2-moxNG)* derived RNA which lacks this splice junction (Figure S6). *Tg(scube2:scube2-moxNG)* embryos were markedly resistant to treatment with this morpholino, validating the *in vivo* functionality of this transgene (Figure S6). *Tg(scube2:scube2-moxNG)* embryos showed broad distributions of Scube2 protein during patterning (Fig. 5H-K). Throughout early patterning, Scube2-moxNeonGreen is visible near ventral cells marked by *Tg(shha:mem-mCherry*), although *scube2* mRNA is expressed largely in the dorsal neuroectoderm at this timepoint (Fig. 5H-I, Movie S5-6). By 24 hpf *Tg(scube2:scube2-moxNG)* fluorescence is found distributed throughout the embryo, although expression from *Tg(scube2:moxNG)* is localized to the dorsal-intermediate neuroectoderm. These data further suggest that Scube2’s long range of effect can be explained by diffusion from secreting cells in the intermediate and dorsal neural tube to the source of Shh in the floor plate and notochord.

### Feedback regulation of Scube2 levels is necessary for pattern scaling

To examine the regulation of Scube2 in size-reduced embryos, we performed size reduction on *Tg(scube2:moxNG; shha:mem-mCherry)* embryos and imaged them at 20 hpf. Unlike other observed patterning genes, *scube2* expression levels did not scale in size-reduced embryos but were instead severely reduced (Fig. 6A-C). This finding is consistent with an expander-repressor-like model of Scube2-Shh. In this regime, inhibition of *scube2* expression in size-reduced embryos would contract Shh signaling, enabling adjustment of Shh signaling for a decreased tissue size (Fig. 6D-E).

**Figure 6.**
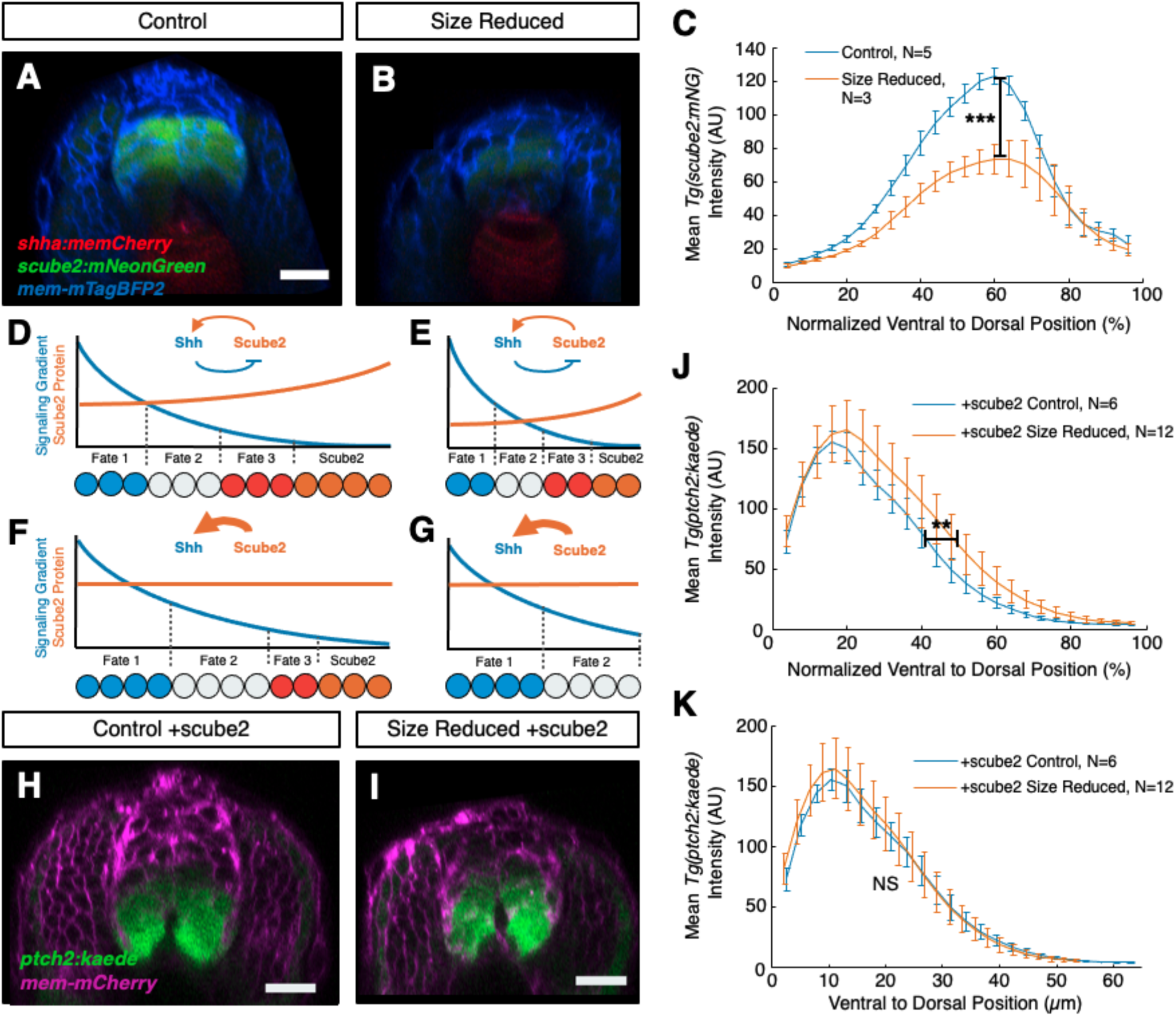
Scube2 expression is size-dependent and required for pattern scaling. (A-B) Transverse view of *mem-mTagBFP2* mRNA-injected *Tg(scube2:moxNG; shha:mem-mCherry)* control (A) or size-reduced (height reduction of 17.1% +/- 7.7%) (B) embryos at 20 hpf. (C) Quantification of mean *Tg(scube2:moxNG)* intensity versus ventral-to-dorsal position of embryos from A-B. Maximum intensity values are statistically significantly reduced in treated embryos (p=3.03e-4). (D-E) Schematic of expander-repressor-like feedback control of Shh signaling by Scube2 and its ability to enable pattern scaling. Repression of Scube2 by Shh encodes an equilibrium level of Shh signaling across the tissue by linking morphogen spread to tissue size. (F-G) Schematic representation of the experiment as performed in H-K, where Scube2 levels are at saturation due to overexpression, and size-reduced embryos (G) are disproportionately affected. (H-I) Transverse view of 20 hpf *Tg(ptch2:kaede)* control (H) and size-reduced (I) embryos injected with *mem-mCherry* and *scube2* mRNA. (J) Quantification of mean *Tg(ptch2:kaede)* intensity versus normalized ventral-to-dorsal position of embryos treated as in H-I. Significant shifts are observed in the dorsal position of 50% of the maximum intensity value (unpaired t-test p=0.0092). (K) Intensity profiles from J rescaled to reflect the absolute scale of the measurements (DV height reduction of 15.2% +/- 2.1%). When measured in these coordinates, no difference is observed in the position of half of maximum intensity (p= 0.9760).

Next, we examined whether feedback control of *scube2* expression levels by Shh signaling is required for pattern scaling. To bypass the feedback regulation, we overexpressed *scube2* at saturation level via mRNA injection into *ptch2:kaede* reporter embryos. If Scube2 is responsible for adjusting Shh signaling during scaling, we would expect *scube2*-overexpressing size-reduced embryos to have the same absolute Shh response profiles as controls, which would fail to scale following size normalization (Fig. 6F-G). If scaling of ventral patterning is not dependent on Scube2, we would expect maintenance of pattern scaling with size-proportionate increases in *ptch2:kaede* distributions in both populations. When normalized for differences in D-V heights, size-reduced *scube2*-overexpressing embryos showed a disproportionate expansion of the *ptch2:kaede* gradient compared to normal-sized *scube2*-overexpressing embryos (Fig. 6H-K). Dorsal expansion of Shh signaling is quantified using the position of 50% of average maximum control intensity, which is statistically significantly shifted dorsally in size-reduced embryos (Fig. 6J). Importantly, when *Tg(ptch2:kaede)* response profiles are plotted on an absolute rather than relative scale, they nearly exactly overlap (Fig. 6K). This overlap without size normalization suggests that *scube2* overexpression encodes a response profile which is independent of embryo size and is not secondarily tuned by another scaling related factor. This strongly suggests that control of *scube2* expression levels is required for scaling the Shh signaling gradient.

### Increase in Shh release and diffusion by Scube2 produces scaling in silico

To examine how Scube2 contributes to scaling, we constructed mathematical models to capture the feedback between Scube2 and Shh, and asked how the biophysical nature of Scube2-Shh protein interaction influences scaling. Like the original expansion-repression model (Ben-Zvi and Barkai, 2010), our experimental results show that the morphogen Shh represses production of Scube2 while Scube2 expands the range of Shh. However, how Scube2 expands the range of Shh remains unclear. In the original expansion-repression model, the expander increases the spread of the morphogen by increasing the diffusion coefficient or decreasing the degradation rate of the morphogen (Ben-Zvi and Barkai, 2010), but does not affect its rate of release. In the case of Scube2, biochemical studies have shown that Scube2 can promote Shh release and Shh is secreted as a complex with Scube2 (Tukachinsky et al., 2012; Wierbowski et al., 2020). However, whether Scube2 increases the diffusion coefficient of Shh or decreases the degradation rate of Shh remains to be experimentally tested. We sought to computationally examine how the Shh and Scube2 feedback circuit would affect scaling when Scube2 acts on 1) release of Shh and 2) release and diffusion of Shh.

We assume that Shh (HH) is released into the patterning field at one boundary at rate κH, diffuses with diffusion constant DH, and is degraded uniformly throughout the field (Fig. 7A). Scube2 (S) is initially produced uniformly throughout the field, but Shh signal activation represses Scube2 production. Scube2 can bind to Shh to form a complex (C), which has possibly modified release rate and diffusion constant, κHS and DHS, respectively. In this generic model, Scube2 can modulate the Shh gradient by increasing its release rate into the patterning field (κHS > κH) and/or promoting its diffusion within the field (DHS > DH). Under assumptions that (1) Scube2 and Shh binding is fast, (2) Scube2 has a fast diffusion constant, and (3) Scube2 concentration is large, Shh concentration becomes an ordinary diffusion first-order degradation equation with a modified diffusion constant and flux (SI Text). To evaluate the scaling capacity of the model, we measure scaling as the average fractional shift in morphogen concentrations at 75%, 50%, and 25% field length, when the field size is reduced from 300 μm to 200 μm (Ben-Zvi and Barkai, 2010). We refer to this as the “domain” scaling metric (ρι), where a smaller ρι corresponds to better scaling (SI Text).

**Figure 7.**
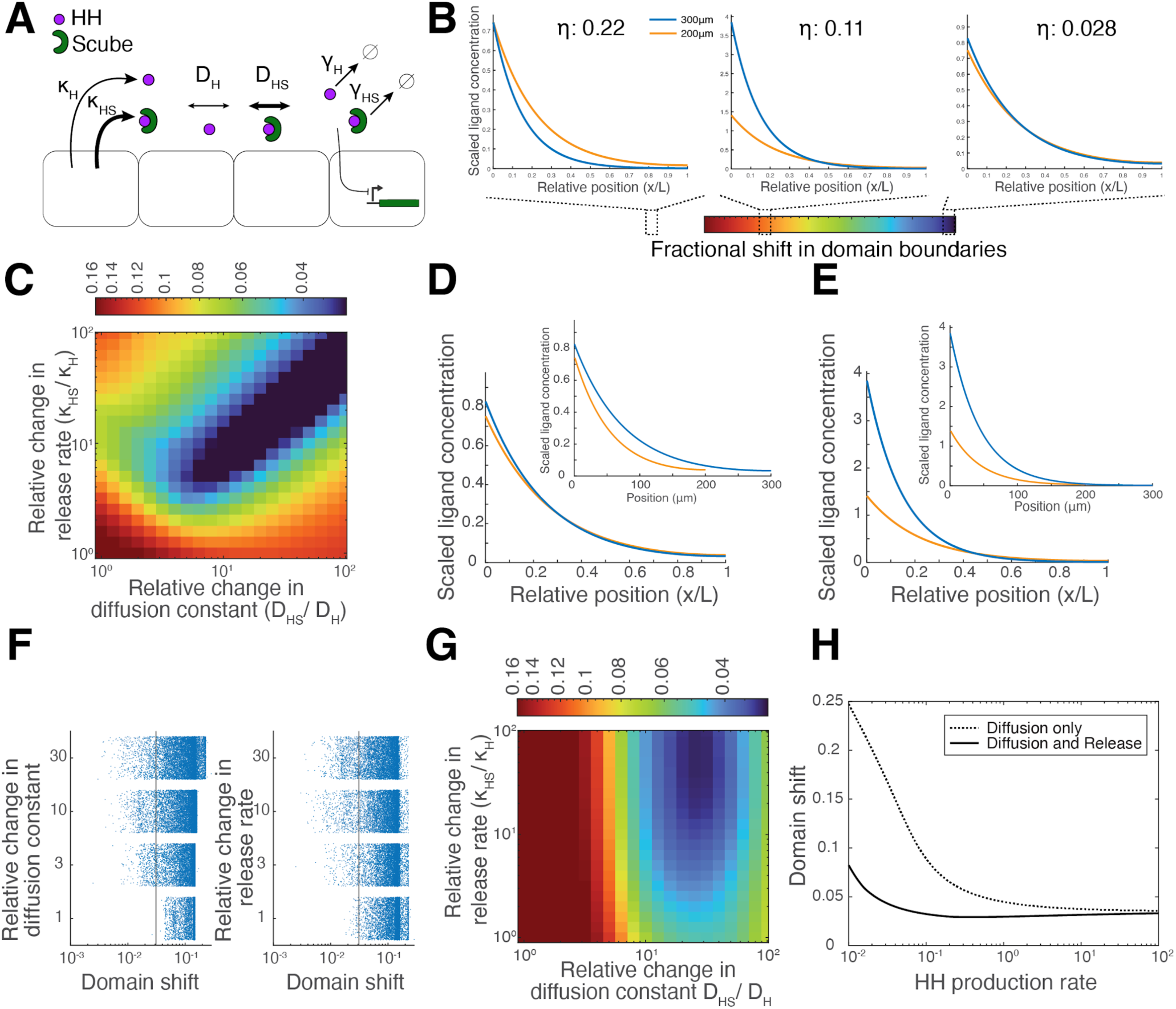
Investigation of effects on scaling by Scube2’s role in release in Hh-Scube feedback. (A) Cartoon of the simplified model representing key interactions. Hedgehog (Hh) in purple and Scube in green. From Hh sending cells, Hh and Hh-Scube complex are released. Thicker arrows indicate that Scube increases the release rate or diffusion constant of Hh. Flux of total Hh (Φ) is dependent on the relative release rates of Hh and of the Hh-Scube complex (κH, κHS), Scube concentration at the source, and the threshold at which Scube affects Hh. In the extracellular space, Hh and Hh-Scube complex diffuse with diffusion rates (DH and DHS, respectively). Hh represses Scube expression, as indicated by the inhibition arrow leading to the Scube-producing cell. Hh and Hh-Scube complex degrade throughout the field at respective rates (γH and γHS). Detailed description and mathematical derivations can be found in supplemental text (SI Text). (B) Example gradient profiles for the scaling metric. Scaling metric (η) is an average of fractional shift in domain boundaries at L3/4, L/2, and L/4. Morphogen profiles and their scaling metrics are shown for 3 example sets of parameters. Orange line indicates ligand concentration (y-axis) by the relative position in the simulated patterning field (x-axis) where patterning field length = 300 μm. Blue line indicates ligand concentration where patterning field length is reduced to 200 μm. (C) Heatmap of scaling metric as a function of relative changes in Hh release rate and diffusion constant, with Hh degradation being uniformly first-order. (D) An example of a set of parameters that produce scaling when Scube affects both release and diffusion of Hh in the uniform degradation model. Orange line indicates ligand concentration (y-axis) by the relative position in the simulated patterning field (x-axis) where patterning field length = 300 μm. Blue line indicates ligand concentration where the patterning field length is reduced to 200 μm. Inset shows the ligand concentration by absolute position from the same simulation. (E) Gradient profile for a model with the same parameters as (D), *except* Scube does not affect diffusion of Hh. As in (D), the orange line indicates ligand concentration where patterning field length = 300 μm and the blue line for field length 200 μm. Inset shows the ligand concentration by absolute position from the same simulation. (F) Plot of parameter sets that produce scaling (x-axis) from a relative change in the diffusion constant (left) or release rate (right) of 1, 3, 10, and 30 (y-axis). Grey line indicates an arbitrary threshold to the left of which are sets of parameters that produce better scaling. (G) Heatmap of scaling metric as a function of relative changes in Hh release rate and diffusion constant, with Hh degradation being self-enhanced. (H) Plot of scaling metric (y-axis) by Hh production rate (x-axis) when Scube affects diffusion only (blue line) and when Scube affects diffusion and release (orange line) in self-enhanced degradation model.

We investigate how the effects of Scube on Shh release and on diffusion influence scaling, first, in a uniform degradation model in which Shh experiences first-order degradation. We ask whether Scube modulation of Shh release alone is sufficient to produce the observed scaling behavior. We found that only models where Scube2 plays both roles (κHS > κH, DHS > DH) could produce scaling behavior as modulation of the amplitude and length scale are required (Fig. 7C-D). In particular, “release-only” models (κHS > κH, DHS = DH) did not scale (Fig. 7C, Fig. 7E). Likewise, “expander-only” models (κHS = κH, DHS > DH) also did not perform as well as models that do both. These results held over a wide range of parameters characterizing Scube2 (Fig. 7F, Fig. S7).

The Hedgehog (Hh) pathway has a unique feedback architecture in which Hh upregulates the expression of its own receptor Patched (Ptch; Chen and Struhl 1996; Goodrich et al., 1996). This evolutionarily conserved feedback architecture provides the system robustness against perturbations in Hh production rate, partly through enhancing ligand degradation and partly through direct suppression of intracellular signaling activity (Goodrich et al., 1997; Denef et al., 2000; Li et al., 2018). Self-enhanced ligand degradation has been implicated in scaling (Eldar et al., 2003; Lander et al., 2009). To ask if self-enhanced ligand degradation could change the requirement for increasing Shh diffusion in the scaling model, we allowed Hh to promote its own degradation. In such models, we found again that a simple increase in Hh release alone was not sufficient to produce scaling (Fig. 7G, Fig. S8). Instead, increasing the diffusion constant could lead to scaling throughout much of the field, but (1) not near the source and (2) only when morphogen release is high enough (ε << L) (SI Text). To capture differences along the entire length of the gradient we also used a “profile” metric which is the mean squared difference between the two gradients at all points (SI Text). Using this more stringent profile scaling metric, Scube2 acting as a releaser in addition to expander allowed modulation of the amplitude which led to better scaling near the source. Further analysis revealed that models where Scube2 increases both diffusion and release led to scaling over a larger set of Hh production rates when compared to models where Scube2 increases only diffusion (Fig. 7H). The requirement for Scube2 to accelerate Shh diffusion in order to scale also remains when considering the self-enhanced degradation feedback mediated by the Hh receptor PTCH (Fig. S8).

Morphogen systems are frequently robust to gene dosage. For example, mouse mutants for one allele for *Shh* are phenotypically wildtype (Chiang et al., 1996). To explore a potential role for the expander-repressor-releasor model in gene dosage robustness, we screened parameter space for models that are robust to halving the Shh production rate. We found that models in which Scube2 enhances the release of Shh promote dosage robustness (Fig. S9, SI Text), which can be in addition to robustness already provided by self-enhanced degradation (Eldar et al., 2003).

In the classic expansion-repression model, the “expander” can also extend the morphogen gradient by repressing the degradation of the morphogen (Ben-Zvi and Barkai, 2010). If Scube2 expands the Hedgehog gradient by decreasing the HH degradation rate, in order to leave the amplitude invariant, Scube2 must also decrease the release rate from the HH secreting cells, which is inconsistent with observed biochemical data (Fig. S10, SI Text). Alternatively, when the size of an animal is decreased, it is possible that the number of ligand-producing cells is correspondingly decreased (“flux scaling”), which could potentially lead to a scaled Hedgehog release rate. However, because the lengthscale of the gradient depends on the diffusion constant and not the rate of Hedgehog flux, the gradient cannot be scaled by simply changing Hedgehog release only (Fig. S11). Taken together, our modeling results suggest that Shh gradient scaling is best achieved by Scube2 both increasing the release of Shh from sender cells, and the diffusion of Shh throughout the field.

## Discussion

Our work uncovers that the morphogen Sonic Hedgehog can self-regulate to enable scale-invariant patterning through linking morphogen signaling to inhibition of Scube2. We discovered that patterning of the neural tube adjusts to tissue availability following surgical size reduction in zebrafish embryos. Using overexpression experiments we demonstrate that Scube2’s activity during patterning is not just permissive—overexpression of *scube2* instead enhances Shh signaling (Woods and Talbot, 2005). Utilizing a transgenic reporter line which we developed, we characterized the expression of Scube2 during neural patterning and found that Shh signaling is responsible for its repression in the ventral neural tube. Using Scube2 fluorescent fusion proteins we found that Scube2 is broadly distributed from secreting cells, explaining its previously reported cell non-autonomous activity (Creanga et al., 2012; Woods and Talbot, 2005). Unlike other patterning genes, *scube2* responds to reduction in neural tube height by disproportionately decreasing its expression, and overexpression of Scube2 inhibits scaling of the Shh signaling gradient by circumventing its feedback control. The expression of *scube2* thus can be seen as a “size-dependent factor”, which can enable scaling by tuning expression levels to embryo size through feedback inhibition (Inomata et al., 2013).

The relationship between Scube2 and Shh has important similarities to proposed “expander-repressor” models of morphogen scaling (Barkai and Ben-Zvi, 2009; Ben-Zvi and Barkai, 2010; Inomata et al., 2013). As with expanders in these models, *scube2* enhances morphogen range, is repressed by morphogen signaling, and acts cell non-autonomously at a distance from its source. However, the reported role of Scube2 in morphogen release is distinct from the proposed mechanism of expanders. Expanders extend the range of morphogens such as by promoting their diffusion or inhibiting their degradation (Ben-Zvi and Barkai, 2010).

We examined the effect of release on scaling through mathematical modeling. Our results show that while increasing release alone does not allow for scaling, Scube2’s role in release allows for modulation of gradient amplitude in uniform and self-enhanced degradation models, where production of Hh is not otherwise coupled to the size of the field, and we find that models in which Scube2 acts as both an expander and releaser perform better. *In vivo*, we showed that decreasing scube2 levels via morpholino knockdown led to a significant decrease in the amplitude of *Tg(ptch2:kaede)* gradient (Fig. 2C), which aligns with Scube2’s effect on gradient amplitude in the modeling. Furthermore, in our model, Scube2’s effect on release in addition to diffusion broadens the range of Shh production rate over which scaling is achieved. However, since we made certain simplifying assumptions and evaluated scaling once equilibrium was reached, there may be other contributions to gradient scaling from the release role of Scube2. In our model, we did not include other pathway features, such as Patched negative regulation of Shh (Chen and Struhl, 1996; Goodrich et al., 1996). In models examining the negative feedback architecture, Patched adds robustness to Shh production rate and allows steady state to be reached more quickly (Li et al., 2018). Additionally, our model does not explore the effects on dynamics of gradient formation. Dessaud et al. showed that signal duration of Shh, in addition to the concentration, impacts downstream transcriptional activity and differentiation (Dessaud et al., 2007). How Scube2’s interaction with Shh affects the temporal features of Shh gradient formation would be an interesting avenue for further study.

While the focus of this work is on how morphogen gradients can be robust to changes in the size of the patterning field, it is also important for morphogen systems to be robust to variation in gene expression such as those caused by noise or gene dosage. Notably, despite being proposed to encode information in their quantitative levels, heterozygous mutants in key morphogens are generally phenotypically normal. Mice heterozygous for Shh are reported to develop normally (Chiang et al., 1996). The situation in zebrafish is more difficult to analyze as there are two paralogs *shha* and *shhb.* We speculate that feedback systems such as the releaser-expander-repressor model presented here allow for robustness to expression levels. Our model supports that feedback on release can give additional robustness to gene dosage on top of that given by self-enhanced degradation (Eldar et al., 2003; Li et al. 2018). Overall, our model shows feedback on morphogen release directly affects gradient amplitude while feedback on morphogen diffusion and degradation primarily affects morphogen length scale. Although, for Scube2, its role in regulating the release and diffusion rate may be biochemically coupled, other morphogen systems may use different regulators to achieve robustness to tissue size.

We began this work in part due to interest in the discrepancy between the area of Scube2 activity in the ventral neural tube and its expression in the dorsal neural tube. Our work with Scube2 fluorescent protein fusions revealed that Scube2 is highly diffusive and is distributed broadly from producing cells (Figure 5). Diffusion of Scube2 from producing cells could easily account for the distance between its expression domain and area of effect. Scube2’s broad distribution and considerable extracellular diffusivity bolsters the hypothesis that Scube2 may serve as a chaperone for Shh during its transport as hypothesized previously (Tukachinsky et al., 2012). Cell culture experiments have indicated that Scube2 cooperates with Dispatched in a cholesterol-dependent “hand off” by binding different domains on Shh’s cholesterol moiety (Tukachinsky et al., 2012). Furthermore, in cell culture, Scube2 complexes with Shh during release via Shh’s lipid modifications, and this lipid-mediated interaction is later dissociated to form Shh-Ptch by coreceptors in the responding cells (Wierbowski et al., 2020). Continued binding of Scube2 to the hydrophobic sterol domain may facilitate the diffusion of Shh through the extracellular millieu. This model is consistent with the dose dependency we observe in our Scube2 overexpression experiments (Figure 2G-L) and may help solve the puzzle of the long-range transport of dually lipid modified hedgehog in vertebrates. Further investigation of the strength and duration of Scube2 and Shh’s binding *in vivo* and Scube2’s effects on diffusion rates of Shh may shed light on this relationship. Unfortunately, direct imaging of this phenomenon is hampered by the lack of fully functional Shh fluorescent protein fusions (Chamberlain et al., 2008).

While some evidence suggests that Scube2 plays a role in lipid-shedding, these observations conflict with previous HPLC analysis and the findings of independent groups which demonstrated that Shh species released by Scube2 are dually lipid modified (Creanga et al., 2012; Tukachinsky et al., 2012). Additionally, Shh and Scube2 were shown to comigrate in native gel electrophoresis (Wierbowski et al., 2020). If correct, a model of Scube2 in which it acts only transiently at the cell surface of producing cells—either by enabling the formation of multimeric Shh complexes or lipid shedding—would have interesting implications for its role as an expander. Expanders are often formalized as having a dose dependent reversible effect on morphogen spread, while a transient role of Scube2 in Shh multimeric complex formation or shedding would be localized and irreversible. In our mathematical modeling, we found that increasing release alone did not produce scaling (Fig. 7C, Fig. 7G), suggesting that there would need to be another mechanism by which lengthscale of Shh could be modulated if Scube2 were to affect release only.

Scube2 is one of several recently identified elements of the Shh signaling pathway that exerts cell non-autonomous effects. Recent work has shown that Hhip—initially characterized as a membrane-tethered Hedgehog antagonist—acts over a long range that cannot be explained by ligand sequestration (Kwong et al., 2014). Additionally, the Hedgehog receptor, Patched, may also have cell non-autonomous inhibitory effects on Smoothened through regulating inhibitory sterols or sterol availability (Bidet et al., 2011; Roberts et al., 2016). Together with known feedback relationships and the diffusivity of Scube2 that we demonstrated here, these mechanisms interlink Shh signaling between neighboring cells and may enable tissue level properties, such as the scaling of pattern formation we observed.

## Methods

### Generation of Transgenic Lines

The construct used to make *Tg(scube2:moxNG)* was generated by isothermal assembly of PCR-amplified *scube2* regulatory elements obtained from the CHORI-211 BAC library. Regulatory elements were in part chosen based on annotations of H3K4me1 and H3K4me3 binding (Aday et al., 2011). Selected regulatory sequences spanned 1677bp of upstream intergenic sequence and 5962bp of the area spanning exons 1-5 *scube2*. Regulatory sequences were cloned into a pMT backbone by placing a zebrafish codon-optimized moxNeonGreen fluorescent protein and sv40 poly-A tail just downstream of the endogenous *scube2* Kozak sequence (Costantini et al., 2015). The construct used to make *Tg(scube2:scube2-moxNG)* was generated using the same regulatory sequences as *Tg(scube2:moxNG*), with the addition of cDNA corresponding to exons 6-23 of the Scube2 coding sequence downstream of exon 5 and moxNeonGreen attached at the c-terminus with a 10 amino acid long GA rich linker. The construct used to make *Tg(shha:mem-mCherry)* was derived from the previously reported *Tg(shh:GFP*), by replacement of GFP with mem-mCherry (Megason, 2009; Shkumatava et al., 2004).

Transgenic lines were generated by injecting plasmid DNA for each construct along with Tol2 mRNA into wild type (AB) embryos at the single cell stage, as described previously (Kawakami, 2004). moxNeonGreen positive embryos were then selected for raising. Upon reaching sexual maturity, F0s were outcrossed and screened for founders. Founders were isolated and raised as single alleles. Monoallelic versions of each line are shown throughout the paper.

### Zebrafish Strains

For wild type lines, AB fish were used. All fish were kept at 28°C on a 14-hour-light/10-hour-dark cycle. Embryos were collected from natural crosses. All fish-related procedures were carried out with the approval of Institutional Animal Care and Use Committee (IACUC) at Harvard University. *TgBAC(ptch2:kaede*) (Huang et al., 2012; renamed from *ptch1* due to a change in zebrafish gene nomenclature), *Tg(nkx2.2a:mGFP)* (Jessen et al., 1998), *Tg(olig2:GFP)* (Shin et al., 2003), *Tg(olig2:dsRed)* (Kucenas et al., 2008), and *TgBAC*(*dbx1b:GFP)* (Kinkhabwala et al., 2011) have been described previously.

### Size Reduction Technique

Size reduction was performed as described in our previous report (Ishimatsu et al., 2018, 2019). Embryo sizes were reduced by sequentially removing ∼1/3 of the cells from the animal cap, then wounding the yolk. These surgeries are performed in 1/3 ringers solution, and embryos are immobilized in a thin layer of 2% methyl cellulose. Surgeries can be performed either with glass needles – as previously described – or using a loop of thin stainless-steel wire that is inserted through a glass capillary tube and mounted on a halved chopstick as done here. Healthy uninjected embryos show a maximum success rate of ∼60% while embryos which have undergone injection or were spawned by older females have significantly lower success rates. In each size reduction experiment, embryos are screened for health and the largest size reductions; those with insufficient size reduction or with morphological defects are discarded.

### Construct Generation and Injections of mRNAs and Morpholinos

Scube2-mCitrine was generated from cDNA obtained from the Talbot lab (Woods and Talbot, 2005). Fluorescent protein fusions were made by attaching mCitrine or moxNeonGreen with a 10 amino acid GA rich linker to the c-terminus of Scube2. Membrane-mTagBFP2 constructs were generated using membrane localization tags reported previously (Megason, 2009; Subach et al., 2011). These constructs were each sub-cloned into a pMTB backbone. mRNA for all experiments was synthesized from pCS or pMTB backbones via *in vitro* transcription using the mMESSAGE mMACHINE system (Ambion). Embryos were injected at the single cell stage using a Nanoject system set to 2.3nl of injection volume containing 70-90pg of RNA for each mRNA injected. Injected embryos were then screened for brightness, and damaged embryos were removed. Scube2 morpholino injections were performed with 7ng of Scube2 MO2 and 3.5ng of p53 MO to control for phenotypic variability, while control morpholino injections were performed using 10.5ng of p53 MO only (Gerety and Wilkinson, 2011; Woods and Talbot, 2005).

### Sonidegib and Cyclopamine Treatment

Stock solution of 1 mM Sonidegib suspended in DMSO was used for treatment as generously given by the lab of Rosalind Segal. Embryos were placed in egg water containing a concentration of 50μM for the treatment condition, and equal parts DMSO were added to the sham control. Cyclopamine was dissolved in 100% ethanol to make 50mM stock solution and was diluted for treatment in egg water to 100 μM. Treatment began at 7 hpf and continued until imaging at 22 hpf.

### Confocal Imaging

For quantitative imaging, embryos were staged and mounted in our previously described dorsal mount (Kimmel et al., 1995; Megason, 2009; Xiong et al., 2013) in egg water with 0.01% tricaine (Western Chemical, Inc.). Embryos were manipulated for proper positioning with hair loops, before gently lowering the coverslip. Embryos were not depressed by the coverslip or impinged by the mold, enabling imaging of their normal proportions. Imaging was performed on embryos staged at 18-24 hpf, unless otherwise noted in corresponding figure legends. Live imaging was performed using a Zeiss 710 confocal/2-photon microscope, Zen image acquisition software, and C-Apochromat 40X 1.2 NA objective. For fluorescent protein excitation, 405 nm (BFP), 488 nm (GFP/moxNeonGreen), 514 nm (mCitrine), 561 nm (mCherry/dsRed) and 594 nm (mCardinal) lasers were used. The imaging field was centered in each embryo on the somite 6/7 boundary for consistent positioning between images. For quantitative analysis, imaging datasets are only compared between sibling embryos imaged on the same day with the same settings. This approach aims to avoid clutch effects or variability in detector sensitivity and laser power that occur over time. Typical imaging settings with the 40x objective were as follows: image size of 1024×1024 pixels with .21μm per pixel and an interval of 1μm in the Z direction. For display purposes, images are rendered in cross sectional views (X-Z axis) which are then rotated for display, with image intensities for co-injection markers adjusted evenly within datasets for displayable brightness. FRAP, early stage embryo imaging and time-lapses were performed using a 1.0 NA 20x objective. Brightfield and widefield fluorescence images of whole embryos were obtained using an Olympus MVX10 and a Leica MZ12.5 dissecting microscope.

### FRAP Experiments and Analysis

Imaging for FRAP was performed using a 1.0 NA 20x objective at dome stage. Bleaching was performed for two minutes in a 100umx100um area in the center of the frame with a 488nm Argon laser. Imaging was performed with a low laser power to reduce bleaching, and images were obtained at 10 second intervals over five minutes to quantify recovery. FRAP data in the bleached region was then normalized to the minimum and maximum intensity for each respective time trace. Normalized recovery intensities were then fitted to the following exponential to determine the mobile fraction (*A*) in matlab: *y* = *A*(1 − *e*^−τ^) (Munjal et al., 2015).

### Image Analysis

Images were analyzed using a custom MATLAB-based image analysis software that enables rapid segmentation of neural tube imaging data. Neural imaging data is segmented by the user sequentially from anterior to posterior. Over a set step size (usually 50 pixels), the user selects points at the base of the floor plate cell and top of the roof plate cell that divide the neural tube into its two halves (Fig. S1A). The user then selects the widest point of the neural tube in each image. Imaging data from mature neurons, found laterally, and within the lumen of the neural tube, found medially, are disregarded using a set percentage of neural width (Fig. S1B). Once these positions are recorded, imaging data is then recovered as average pixel intensity in 25 bins from ventral to dorsal across 3-4 somites of A-P length. This binning and averaging strategy enables comparison of data between embryos that accounts for variations in neural tube D-V height. During the segmentation process, the researcher is blinded to the title of the dataset which contains information about its treatment condition. For distribution plots, binned intensities are reported for each embryo as the average intensities for each bin along the entire AP axis of the imaging volume. Each embryo’s average intensity profile is then treated as an independent sample and averaged for displayed distribution profiles and standard deviations. To avoid artifacts caused by rounding in the calculations of half maximum control intensity positional values are extracted from the spline-interpolated intensity profiles for each individual dataset.

Progenitor domain segmentation is performed on average intensity profiles from each embryo in a dataset in the following manner: first, all intensity profiles in the data set undergo background subtraction and cross channel fluorescence caused by the extreme brightness of the *dbx1b:GFP* line is removed from the *olig2:dsred* channel. Intensity profiles are then fed to a peak finding algorithm to identify local maxima. Both *dbx1b+* and *nkx2.2a+* progenitor domains are found in the green channel, so a maximum of two peaks is allowed. In the red channel, only one peak is specified to identify *olig2:dsred* signal. Average peak intensity values for each domain are then calculated for the entire control dataset, and 50% of this value in the case of the *nkx2.2a* and *dbx1b* domains is used as the threshold for calculating domain width. Given its greater spread along the D-V axis, a threshold of 25% of peak height is used in calculating width of the *olig2+* domain. Domain widths are then extracted from spline-interpolated intensity profiles to avoid errors introduced by rounding to the next bin. Segmented widths and positions of *nkx2.2a*, *olig2*, and *dbx1b* expression are then averaged for plotting purposes. Domain plots are generated by assigning all *nkx2.2a*+ progenitors to the p3 fate, *olig2*+ progenitors lacking *nkx2.2a* expression overlap to the pMN fate, and *dbx1b*+ progenitors to the p0-d6 fate. These domain sizes and positions are then used to reconstruct domains in-between or flanking them, which include the p2-p1 domain between pMN and p0-d6, the floorplate below p3, and the d5-roofplate above p0-d6. These heights and positions are then used to generate the stacked bar plots shown.

### Statistical Analysis

Statistical comparisons of maximum average intensity and position of 50% maximum intensity are performed by an unpaired T-test. Although each dataset contains hundreds of measurements of each binned intensity value over the A-P axis of a z-stack, only the average of these measurements for each embryo is treated as a data-point for calculation of the standard deviation and statistical significance tests. This is done to avoid oversampling that would exaggerate statistical significance. In all measurements, statistical significance is markedly increased if analysis is performed by treating all underlying intensity measurements as samples. Thresholds for calculating the position of half maximum are determined from the average maximum of the corresponding control dataset for each experiment. Position is then determined from the fitted trend-line to avoid inaccuracies due to rounding. To calculate the significance of shifts in boundary positions, upper domain boundaries for each embryo were compared in an unpaired t-test between embryos from each population. When the progenitor domain segmentation algorithm finds there is no domain present, the boundary is set to 0.

### CRISPR Screen for Scube2 Regulators

Cas9 protein was generated and purified in lab as described (Gagnon et al., 2014). Three guide RNA sequences targeting the first one-to-three exons of each gene were selected based on their quality using the web-tool CHOP-CHOP and synthesized using standard methods (Gagnon et al., 2014). Equivalent guide RNA and Cas9 protein concentrations were used in all samples for mosaic knockout. Phenotypes were assessed at 18-20 hpf by confocal microscopy.

### Mathematical modeling

See SI Text for full details.

## Acknowledgments

We thank Lisa Goodrich, Rosalind Segal, and Wolfram Goessling for their comments and helpful discussion. The Scube2 construct was a gift from the Talbot lab.

## References

Aday, A. W., Zhu, L. J., Lakshmanan, A., Wang, J. and Lawson, N. D. (2011). Identification of cis regulatory features in the embryonic zebrafish genome through large-scale profiling of H3K4me1 and H3K4me3 binding sites. Dev. Biol. 357, 450–462.

Averbukh I, Ben-Zvi D, Mishra S, Barkai N, Averbukh, I., Ben-Zvi, D., Mishra, S. and Barkai, N. (2014). Scaling morphogen gradients during tissue growth by a cell division rule. Development 141, 2150–6.

Barkai, N. and Ben-Zvi, D. (2009). “Big frog, small frog”--maintaining proportions in embryonic development: delivered on 2 July 2008 at the 33rd FEBS Congress in Athens, Greece. FEBS J. 276, 1196–207.

Ben-Zvi, D. and Barkai, N. (2010). Scaling of morphogen gradients by an expansion-repression integral feedback control. Proc. Natl. Acad. Sci. U. S. A. 107, 6924–9.

Ben-Zvi, D., Shilo, B.-Z., Fainsod, A. and Barkai, N. (2008). Scaling of the BMP activation gradient in Xenopus embryos. Nature 453, 1205–11.

Ben-Zvi, D., Pyrowolakis, G., Barkai, N. and Shilo, B.-Z. (2011). Expansion-repression mechanism for scaling the Dpp activation gradient in Drosophila wing imaginal discs. Curr. Biol. 21, 1391–6.

Ben-Zvi, D., Fainsod, A., Shilo, B.-Z. and Barkai, N. (2014). Scaling of dorsal-ventral patterning in the Xenopus laevis embryo. Bioessays 36, 151–6.

Bidet, M., Joubert, O., Lacombe, B., Ciantar, M., Nehmé, R., Mollat, P., Brétillon, L., Faure, H., Bittman, R., Ruat, M., et al. (2011). The Hedgehog Receptor Patched Is Involved in Cholesterol Transport. PLoS One 6, e23834.

Briscoe, J. and Small, S. (2015). Morphogen rules: design principles of gradient-mediated embryo patterning. Development 142, 3996–4009.

Briscoe, J., Chen, Y., Jessell, T. M. and Struhl, G. (2001). A hedgehog-insensitive form of patched provides evidence for direct long-range morphogen activity of sonic hedgehog in the neural tube. Mol. Cell 7, 1279–91.

Burke, R., Nellen, D., Bellotto, M., Hafen, E., Senti, K.-A. A., Dickson, B. J. and Basler, K. (1999). Dispatched, a Novel Sterol-Sensing Domain Protein Dedicated to the Release of Cholesterol-Modified Hedgehog from Signaling Cells. Cell 99, 803–815.

Cao, Y., Ryser, M. D., Payne, S., Li, B., Rao, C. V. and You, L. (2016). Collective Space-Sensing Coordinates Pattern Scaling in Engineered Bacteria. Cell 165, 620–630.

Čapek, D., & Müller, P. (2019). Positional information and tissue scaling during development and regeneration. Development (Cambridge, England), 146(24), dev177709. https://doi.org/10.1242/dev.177709

Chamberlain, C. E., Jeong, J., Guo, C., Allen, B. L. and McMahon, A. P. (2008). Notochord-derived Shh concentrates in close association with the apically positioned basal body in neural target cells and forms a dynamic gradient during neural patterning. Development 135, 1097–106.

Chen, M.-H., Li, Y.-J., Kawakami, T., Xu, S.-M. and Chuang, P.-T. (2004). Palmitoylation is required for the production of a soluble multimeric Hedgehog protein complex and long-range signaling in vertebrates. Genes Dev. 18, 641–59.

Chen, Y. and Struhl, G. (1996). Dual roles for patched in sequestering and transducing Hedgehog. Cell 87, 553–563. doi: 10.1016/s0092-8674(00)81374-4

Chiang, C., Litingtung, Y., Lee, E., Young, K. E., Corden, J. L., Westphal, H., & Beachy, P. A. (1996). Cyclopia and defective axial patterning in mice lacking Sonic hedgehog gene function. Nature, 383(6599), 407–413. https://doi.org/10.1038/383407a0

Cooke, J. (1981). Scale of body pattern adjusts to available cell number in amphibian embryos. Nature 290, 775–778.

Costantini, L. M., Baloban, M., Markwardt, M. L., Rizzo, M., Guo, F., Verkhusha, V. V. and Snapp, E. L. (2015). A palette of fluorescent proteins optimized for diverse cellular environments. Nat. Commun. 6, 7670.

Creanga, A., Glenn, T. D., Mann, R. K., Saunders, A. M., Talbot, W. S. and Beachy, P. A. (2012). Scube/You activity mediates release of dually lipid-modified Hedgehog signal in soluble form. Genes Dev. 26, 1312–25.

Denef, N., Neubuser, D., Perez, L., and Cohen, S.M. (2000). Hedgehog induces opposite changes in turnover and subcellular localization of patched and smoothened. Cell 102, 521–531. doi: 10.1016/s0092-8674(00)00056-8

Dessaud, E., Yang, L., Hill, K., Cox, B., Uloa, F., Ribeiro, A., Mynett, A., Novitch, B.G. and Briscoe, J., (2007). Interpretation of the sonic hedgehog morphogen gradient by a temporal adaptation mechanism. Nature 450, 717–720. doi:10.1038/nature06347

Driesch, H. (1892). Entwicklungsmechanische Studien: I. Der Werthe der beiden ersten Furchungszellen in der Echinogdermenentwicklung. Experimentelle Erzeugung von Theil-und Doppelbildungen. Zeitschrift für wissenschaftliche Zool.

Eldar, A., Rosin, D., Shilo, B.-Z. and Barkai, N. (2003). Self-Enhanced Ligand Degradation Underlies Robustness of Morphogen Gradients. Dev Cell 5, 635–646. doi: 10.1016/s1534-5807(03)00292-2

Francois, P., Vonica, A., Brivanlou, A. H. and Siggia, E. D. (2009). Scaling of BMP gradients in Xenopus embryos. Nature 461, E1–E1.

Gagnon, J. A., Valen, E., Thyme, S. B., Huang, P., Ahkmetova, L., Pauli, A., Montague, T. G., Zimmerman, S., Richter, C., Schier, A. F., et al. (2014). Efficient Mutagenesis by Cas9 Protein-Mediated Oligonucleotide Insertion and Large-Scale Assessment of Single-Guide RNAs. PLoS One 9, e98186.

Gerety, S. S. and Wilkinson, D. G. (2011). Morpholino artifacts provide pitfalls and reveal a novel role for pro-apoptotic genes in hindbrain boundary development. Dev. Biol. 350, 279–89.

Gribble, S. L., Nikolaus, O. B. and Dorsky, R. I. (2007). Regulation and function of Dbx genes in the zebrafish spinal cord. Dev. Dyn. 236, 3472–83.

Goodrich, L.V., Johnson, R.L., Milenković, L., McMahon, J.A. and Scott, M.P. (1996). Conservation of the hedgehog/patched signaling pathway from flies to mice: induction of a mouse patched gene by Hedgehog. Genes Dev 10, 301–312. doi: 10.1101/gad.10.3.301

Goodrich, L.V., Milenković, L., Higgins, K.M. and Scott, M.P. (1997). Altered Neural Cell Fates and Medulloblastoma in Mouse patched Mutants. Science 277, 1109–1113. doi: 10.1126/science.277.5329.1109

Grimmond, S., Larder, R., Van Hateren, N., Siggers, P., Morse, S., Hacker, T., Arkell, R. and Greenfield, A. (2001). Expression of a novel mammalian epidermal growth factor-related gene during mouse neural development. Elsevier.

Hamaratoglu, F., de Lachapelle, A. M., Pyrowolakis, G., Bergmann, S. and Affolter, M. (2011). Dpp signaling activity requires Pentagone to scale with tissue size in the growing Drosophila wing imaginal disc. PLoS Biol. 9, e1001182.

Hammerschmidt, M., Bitgood, M. J. and McMahon, A. P. (1996). Protein kinase A is a common negative regulator of Hedgehog signaling in the vertebrate embryo. Genes Dev. 10, 647–58.

Hollway, G. E., Maule, J., Gautier, P., Evans, T. M., Keenan, D. G., Lohs, C., Fischer, D., Wicking, C. and Currie, P. D. (2006). Scube2 mediates Hedgehog signalling in the zebrafish embryo. Dev. Biol. 294, 104–18.

Huang, P., Xiong, F., Megason, S. G., & Schier, A. F. (2012). Attenuation of Notch and Hedgehog signaling is required for fate specification in the spinal cord. PLoS genetics, 8(6), e1002762. https://doi.org/10.1371/journal.pgen.1002762

Inomata, H., Shibata, T., Haraguchi, T. and Sasai, Y. (2013). Scaling of dorsal-ventral patterning by embryo size-dependent degradation of Spemann’s organizer signals. Cell 153, 1296–311.

Ishimatsu, K., Hiscock, T. W., Collins, Z. M., Sari, D. W. K., Lischer, K., Richmond, D. L., Bessho, Y., Matsui, T. and Megason, S. G. (2018). Size-reduced embryos reveal a gradient scaling-based mechanism for zebrafish somite formation. Development 145, dev161257.

Ishimatsu, K., Cha, A., Collins, Z. M., & Megason, S. G. (2019). Surgical Size Reduction of Zebrafish for the Study of Embryonic Pattern Scaling. Journal of visualized experiments: JoVE, (147), 10.3791/59434. https://doi.org/10.3791/59434

Jakobs, P., Exner, S., Schürmann, S., Pickhinke, U., Bandari, S., Ortmann, C., Kupich, S., Schulz, P., Hansen, U., Seidler, D. G., et al. (2014). Scube2 enhances proteolytic Shh processing from the surface of Shh-producing cells. J. Cell Sci. 127, 1726–37.

Jakobs, P., Schulz, P., Ortmann, C., Schürmann, S., Exner, S., Rebollido-Rios, R., Dreier, R., Seidler, D. G. and Grobe, K. (2016). Bridging the gap: heparan sulfate and Scube2 assemble Sonic hedgehog release complexes at the surface of producing cells. Sci. Rep. 6, 26435.

Jessen, J. R., Meng, A., McFarlane, R. J., Paw, B. H., Zon, L. I., Smith, G. R. and Lin, S. (1998). Modification of bacterial artificial chromosomes through Chi-stimulated homologous recombination and its application in zebrafish transgenesis. Proc. Natl. Acad. Sci. 95, 5121–5126.

Kawakami, K. (2004). Transgenesis and Gene Trap Methods in Zebrafish by Using the Tol2 Transposable Element.pp. 201–222.

Kawakami, T., Kawcak, T., Li, Y.-J., Zhang, W., Hu, Y. and Chuang, P.-T. (2002). Mouse dispatched mutants fail to distribute hedgehog proteins and are defective in hedgehog signaling. Development 129, 5753–65.

Kawakami, A., Nojima, Y., Toyoda, A., Takahoko, M., Satoh, M., Tanaka, H., Wada, H., Masai, I., Terasaki, H., Sakaki, Y., et al. (2005). The zebrafish-secreted matrix protein you/scube2 is implicated in long-range regulation of hedgehog signaling. Curr. Biol. 15, 480–8.

Kicheva, A., Bollenbach, T., Ribeiro, A., Valle, H. P., Lovell-Badge, R., Episkopou, V. and Briscoe, J. (2014). Coordination of progenitor specification and growth in mouse and chick spinal cord. Science (80-.). 345, 1254927–1254927.

Kimmel, C. B., Ballard, W. W., Kimmel, S. R., Ullmann, B. and Schilling, T. F. (1995). Stages of embryonic development of the zebrafish. Dev. Dyn. 203, 253–310.

Kinkhabwala, A., Riley, M., Koyama, M., Monen, J., Satou, C., Kimura, Y., Higashijima, S.-I. and Fetcho, J. (2011). A structural and functional ground plan for neurons in the hindbrain of zebrafish. Proc. Natl. Acad. Sci. U. S. A. 108, 1164–9.

Kucenas, S., Takada, N., Park, H.-C., Woodruff, E., Broadie, K. and Appel, B. (2008). CNS-derived glia ensheath peripheral nerves and mediate motor root development. Nat. Neurosci. 11, 143–151.

Kwong, L., Bijlsma, M. F. and Roelink, H. (2014). Shh-mediated degradation of Hhip allows cell autonomous and non-cell autonomous Shh signalling. Nat. Commun. 5, 4849.

Lander, A. D., Lo, W. C., Nie, Q., & Wan, F. Y. (2009). The measure of success: constraints, objectives, and tradeoffs in morphogen-mediated patterning. Cold Spring Harbor perspectives in biology, 1(1), a002022. https://doi.org/10.1101/cshperspect.a002022

Li, P., Markson, J. S., Wang, S., Chen, S., Vachharajani, V., & Elowitz, M. B. (2018). Morphogen gradient reconstitution reveals Hedgehog pathway design principles. Science (New York, N.Y.), 360(6388), 543–548. https://doi.org/10.1126/science.aao0645

Liem, K. F., Jessell, T. M. and Briscoe, J. (2000). Regulation of the neural patterning activity of sonic hedgehog by secreted BMP inhibitors expressed by notochord and somites. Development 127, 4855–4866.

Liu, T.-L., Upadhyayula, S., Milkie, D. E., Singh, V., Wang, K., Swinburne, I. A., Mosaliganti, K. R., Collins, Z. M., Hiscock, T. W., Shea, J., et al. (2018). Observing the cell in its native state: Imaging subcellular dynamics in multicellular organisms. Science 360, eaaq1392.

Martí E, Bumcrot DA, Takada R, McMahon AP. (1995). Requirement of 19K form of Sonic hedgehog for induction of distinct ventral cell types in CNS explants. Nature 375, 322–5.

McHale, P., Rappel, W.-J. and Levine, H. (2006). Embryonic pattern scaling achieved by oppositely directed morphogen gradients. Phys. Biol. 3, 107–120.

Megason, S. G. (2009). In toto imaging of embryogenesis with confocal time-lapse microscopy. Methods Mol. Biol. 546, 317–32.

Morgan, T. H. (1895). Half embryos and whole embryos from one of the first two blastomeres. Anat. Anz. 10, 623–638.

Munjal, A., Philippe, J.-M., Munro, E. and Lecuit, T. (2015). A self-organized biomechanical network drives shape changes during tissue morphogenesis. Nature 524, 351–355.

Pepinsky, R. B., Zeng, C., Wen, D., Rayhorn, P., Baker, D. P., Williams, K. P., Bixler, S. A., Ambrose, C. M., Garber, E. A., Miatkowski, K., et al. (1998). Identification of a Palmitic Acid-modified Form of Human Sonic hedgehog. J. Biol. Chem. 273, 14037–14045.

Petrov, K., Wierbowski, B.M. and Salic, A. (2017) Sending and Receiving Hedgehog Signals. Annu.Rev. Cell Dev. Biol. 33, 145–168. doi: 10.1146/annurev-cellbio-100616-060847

Pierani, A., Brenner-Morton, S., Chiang, C. and Jessell, T. M. (1999). A sonic hedgehog-independent, retinoid-activated pathway of neurogenesis in the ventral spinal cord. Cell 97, 903– 15.

Porter, J. A., Ekker, S. C., Park, W.-J., von Kessler, D. P., Young, K. E., Chen, C.-H., Ma, Y., Woods, A. S., Cotter, R. J., Koonin, E. V, et al. (1996a). Hedgehog Patterning Activity: Role of a Lipophilic Modification Mediated by the Carboxy-Terminal Autoprocessing Domain. Cell 86, 21– 34.

Porter, J. A., Young, K. E., & Beachy, P. A. (1996b). Cholesterol modification of hedgehog signaling proteins in animal development. Science (New York, N.Y.), 274(5285), 255–259. https://doi.org/10.1126/science.274.5285.255

Roberts, B., Casillas, C., Alfaro, A. C., Jägers, C. and Roelink, H. (2016). Patched1 and Patched2 inhibit Smoothened non-cell autonomously. Elife 5,.

Roelink H, Porter JA, Chiang C, Tanabe Y, Chang DT, Beachy PA, Jessell TM. (1995). Floor plate and motor neuron induction by different concentrations of the amino-terminal cleavage product of sonic hedgehog autoproteolysis. Cell 81, 445–55.

Romanova-Michaelides, M., Hadjivasiliou, Z., Aguilar-Hidalgo, D., Basagiannis, D., Seum, C., Dubois, M., Jülicher, F., & Gonzalez-Gaitan, M. (2022). Morphogen gradient scaling by recycling of intracellular Dpp. Nature, 602(7896), 287–293. https://doi.org/10.1038/s41586-021-04346-w

Sanders, T.A., Llagostera, E. and Barna, M. (2013). Specialized filopodia direct long-range transport of SHH during vertebrate tissue patterning. Nature 497, 628–632. doi: 10.1038/nature12157.

Shilo, B.-Z. and Barkai, N. (2017). Buffering Global Variability of Morphogen Gradients. Dev. Cell 40, 429–438.

Shin, J., Park, H.-C., Topczewska, J. M., Mawdsley, D. J. and Appel, B. (2003). Neural cell fate analysis in zebrafish using olig2 BAC transgenics. Methods Cell Sci. 25, 7–14.

Shkumatava, A., Fischer, S., Müller, F., Strahle, U. and Neumann, C. J. (2004). Sonic hedgehog, secreted by amacrine cells, acts as a short-range signal to direct differentiation and lamination in the zebrafish retina. Development 131, 3849–58.

Spemann, H. (1938). Embryonic Development and Induction. Yale Univ.; New Haven:

Subach, O. M., Cranfill, P. J., Davidson, M. W. and Verkhusha, V. V. (2011). An Enhanced Monomeric Blue Fluorescent Protein with the High Chemical Stability of the Chromophore. PLoS One 6, e28674.

Tuazon FB, Wang X, Andrade JL, Umulis D, Mullins MC. (2020). Proteolytic Restriction of Chordin Range Underlies BMP Gradient Formation. Cell Rep. 32, 108039.

Tukachinsky, H., Kuzmickas, R. P. P., Jao, C. Y. Y., Liu, J. and Salic, A. (2012). Dispatched and Scube Mediate the Efficient Secretion of the Cholesterol-Modified Hedgehog Ligand. Cell Rep. 2, 308–320.

Umulis, D. M. and Othmer, H. G. (2013). Mechanisms of scaling in pattern formation. Development 140, 4830–43.

Uygur, A., Young, J., Huycke, T. R., Koska, M., Briscoe, J. and Tabin, C. J. (2016). Scaling Pattern to Variations in Size during Development of the Vertebrate Neural Tube. Dev. Cell 37, 127–135.

van Eeden, F. J. J., Granato, M., Schach, U., Brand, M., Furutani-Seiki, M., Haffter, P., Hammerschmidt, M., Heisenberg, C. P. P., Jiang, Y. J. J., Kane, D. A. A., et al. (1996). Mutations affecting somite formation and patterning in the zebrafish, Danio rerio. Development 123, 153–164.

Wierbowski, B. M., Petrov, K., Aravena, L., Gu, G., Xu, Y., & Salic, A. (2020). Hedgehog Pathway Activation Requires Coreceptor-Catalyzed, Lipid-Dependent Relay of the Sonic Hedgehog Ligand. Developmental cell, 55(4), 450–467.e8. https://doi.org/10.1016/j.devcel.2020.09.017

Willot, V., Mathieu, J., Lu, Y., Schmid, B., Sidi, S., Yan, Y.-L., Postlethwait, J. H., Mullins, M., Rosa, F. and Peyriéras, N. (2002). Cooperative Action of ADMP- and BMP-Mediated Pathways in Regulating Cell Fates in the Zebrafish Gastrula. Dev. Biol. 241, 59–78.

Wolpert L. (1969). Positional information and the spatial pattern of cellular differentiation. Journal of theoretical biology, 25(1), 1–47. https://doi.org/10.1016/s0022-5193(69)80016-0

Woods, I. G. and Talbot, W. S. (2005). The you gene encodes an EGF-CUB protein essential for Hedgehog signaling in zebrafish. PLoS Biol. 3, e66.

Xiong, F., Tentner, A. R., Huang, P., Gelas, A., Mosaliganti, K. R., Souhait, L., Rannou, N., Swinburne, I. a, Obholzer, N. D., Cowgill, P. D., et al. (2013). Specified neural progenitors sort to form sharp domains after noisy Shh signaling. Cell 153, 550–61.

Xu, J., Srinivas, B. P., Tay, S. Y., Mak, A., Yu, X., Lee, S. G. P., Yang, H., Govindarajan, K. R., Leong, B., Bourque, G., et al. (2006). Genomewide expression profiling in the zebrafish embryo identifies target genes regulated by Hedgehog signaling during vertebrate development. Genetics 174, 735–52.

Yang, R.-B., Ng, C. K. D., Wasserman, S. M., Colman, S. D., Shenoy, S., Mehraban, F., Komuves, L. G., Tomlinson, J. E. and Topper, J. N. (2002). Identification of a novel family of cell-surface proteins expressed in human vascular endothelium. J. Biol. Chem. 277, 46364–73.

Zagorski, M., Tabata, Y., Brandenberg, N., Lutolf, M. P., Tkačik, G., Bollenbach, T., Briscoe, J. and Kicheva, A. (2017). Decoding of position in the developing neural tube from antiparallel morphogen gradients. Science 356, 1379–1383.

Zeng, X., Goetz, J. A., Suber, L. M., Scott, W. J., Schreiner, C. M. and Robbins, D. J. (2001). A freely diffusible form of Sonic hedgehog mediates long-range signalling. Nature 411, 716–20.

Zhu, Y., Qiu, Y., Chen, W., Nie, Q., & Lander, A. D. (2020). Scaling a Dpp Morphogen Gradient through Feedback Control of Receptors and Co-receptors. Developmental cell, 53(6), 724– 739.e14. https://doi.org/10.1016/j.devcel.2020.05.029

Zinski, J., Bu, Y., Wang, X., Dou, W., Umulis, D., & Mullins, M. C. (2017). Systems biology derived source-sink mechanism of BMP gradient formation. eLife, 6, e22199. https://doi.org/10.7554/eLife.22199

