## Supplement for "A Scube2-Shh feedback loop links morphogen release and spread to morphogen signaling to enable scale invariant patterning of the ventral neural tube"

#### Contents


page

Supplementary Text 1-11
- 1 Hedgehog-Scube model 1
- 2 Modeling setup 4
- 3 Uniform degradation 5
- 4 Self-enhanced degradation 7
- 5 Heterozygous HH knockout 9
- 6 Scube modulating HH degradation 9
- 7 HH flux scales with patterning field size 10

Supplementary Figures S1-S11 11-30
- Figure S1- Segmentation of neural imaging data.
- Figure S2- Progenitor domain width determination and bar plot generation.
- Figure S3- Cyclopamine treatment of tg(scube2:moxNG) embryos.
- Figure S4- Mosaic pax6a/b CRISPR mutants have lowered scube2 expression.
- Figure S5- Rescue of scube2 CRISPR mutants with scube2 or scube2-mCitrine mRNA.
- Figure S6- Rescue of morpholino knockdown by tg(scube2:scube2-moxNeonGreen) expression.
- Figure S7- Uniform degradation model.
- Figure S8- Self-enhanced degradation model.
- Figure S9- Effect of 50% reduction in Hedgehog flux in uniform degradation and self-enhanced degradation models
- Figure S10- Model in which Scube2 enhances degradation of Hedgehog.
- Figure S11- Model in which Hedgehog flux scales with the size of the patterning field.

.

Supplementary Movie Captions S1-S6 31

### Scube-mediated Scaling in the Hedgehog System

#### 1 Hedgehog-Scube model

##### 1.1 Hedgehog production, degradation, and diffusion

We begin with a simple model that captures the production of Hedgehog at the proximal end of the patterning field and the first-order degradation of Hedgehog throughout the patterning field. The Hedgehog concentration in the field $H(x)$ obeys the diffusion equation:

$$\frac{\partial H}{\partial t} = D_H \nabla^2 H - \gamma_H H \tag{1}$$

At the source cells, Hedgehog is produced and accumulates on the cell membrane, where it is then released into the patterning field at some rate $\kappa_H$. We assume that the release of Hedgehog molecules from the membrane is the slow step. Then the concentration of Hedgehog molecules on the surface of the source cells is approximately constant, and the Hedgehog flux into the patterning field is $\Phi_0 = \kappa_H H_0$, where $H_0$ is the Hedgehog concentration on the membrane.

##### 1.2 Phenomenological model of Hedgehog-Scube interaction

We now introduce a new species, Scube. Scube can diffuse with diffusion constant $D_S$, is degraded throughout the field at rate $\gamma_S$, and is produced throughout the field under negative feedback control by Hedgehog (at maximum rate $\alpha_S$ with hill coefficient $n_H$ and repression threshold $K_H$). The Scube concentration in the field $S(x)$ obeys:

$$\frac{\partial S}{\partial t} = D_S \nabla^2 S + \alpha_S \frac{K_H^{n_H}}{K_H^{n_H} + H^{n_H}} - \gamma_S S \tag{2}$$

Scube can bind to Hedgehog and increase its release rate off the cell membrane to some $\kappa_{HS}$. Additionally, Scube may be able to bind Hedgehog and change its diffusive properties within the patterning field. We model these two processes "phenomenologically" - we assume that Scube modifies the local Hedgehog diffusion constant $D(S)$ and Hedgehog flux $\Phi$ in the following way:

$$D(S) = D_H \left[1 + \left(\frac{D_{HS}}{D_H} - 1\right) \frac{S}{K_D + S}\right] \tag{3}$$

$$\Phi = \Phi_0 \left[ 1 + \left( \frac{\kappa_{HS}}{\kappa_H} - 1 \right) \frac{S(0)}{K_D + S(0)} \right] \tag{4}$$

The effect of Scube on Hedgehog is parameterized by the constants $D_{HS}/D_H$, $\kappa_{HS}/\kappa_H$, and $K_D$. $D_{HS}/D_H$ characterizes the effect of Scube on Hedgehog diffusion - it is greater than one for systems where Scube promotes Hedgehog diffusion. $\kappa_{HS}/\kappa_H$ characterizes the effect of Scube on Hedgehog release - it is greater than one for systems where Scube promotes Hedgehog release. $K_D$ characterizes the threshold at which Scube affects Hedgehog.

##### 1.3 Mechanistic model of Hedgehog-Scube interaction

We will now consider a mechanistic model taking into account the binding of Scube and Hedgehog to form a complex with modified release and diffusion properties.

We assume that Scube and Hedgehog reversibly bind to form a Scube-Hedgehog complex ($C$) with rates $k_{on,HS}$ and $k_{off,HS}$. This complex has a (possibly) modified diffusion constant $D_{HS}$ and a (possibly) modified release rate $\kappa_{HS}$. We also assume that Hedgehog is removed from the system primarily via PTCH-mediated endocytosis; then when a Hedgehog molecule comes off the Scube-Hedgehog complex to bind to PTCH and form a PTCH-Hedgehog complex, a Scube molecule is released. Throughout the patterning field, Scube $S$, Hedgehog $H$, and complex $C$ obey the equations:

$$\frac{\partial H}{\partial t} = D_H \nabla^2 H - \gamma_H H + k_{off} C - k_{on} HS \tag{5}$$

$$\frac{\partial S}{\partial t} = D_S \nabla^2 S + \alpha_S \frac{K_H^{n_H}}{K_H^{n_H} + (H+C)^{n_H}} + k_{off} C - k_{on} HS + \gamma_H C - \gamma_S S \tag{6}$$

$$\frac{\partial C}{\partial t} = D_{HS} \nabla^2 C - \gamma_H C - k_{off} C + k_{on} HS \tag{7}$$

We now show that under certain assumptions, the above equations reduce to the phenomenological equations (3) and (4).

We make the following assumptions:

1. The binding kinetics of the Scube-Hedgehog complex are fast: $k_{off} \gg \gamma_H$
2. Scube diffusion is fast: $D_S \gg \alpha_S L^2 / K_D$
3. Scube concentration is large: $\alpha_S \gg \gamma_H H D_S / D_{HS}$

From (1), Hedgehog, Scube, and Hedgehog-Scube complex are in fast equilibrium throughout the patterning field. Then the Hedgehog and Scube concentrations throughout the field satisfy:

$$H \approx \frac{K_D}{S + K_D} H_T \tag{8}$$

where $H_T$ is the *total* Hedgehog concentration and $K_D = k_{off,HS}/k_{on,HS}$. Adding equations (5) and (7) yields a differential equation for total Hedgehog:

$$\frac{\partial H_T}{\partial t} = D_H \nabla^2 H + D_{HS} \nabla^2 C - \gamma_H H_T \tag{9}$$

From assumption (2), the Scube concentration throughout the field is approximately constant. Then, spatial derivatives of $H$ and $C$ are constant multiples of spatial derivatives of $H_T$:

$$\nabla^2 H \approx \frac{K_D}{S + K_D} \nabla^2 H_T \quad \nabla^2 C \approx \frac{S}{S + K_D} \nabla^2 H_T \tag{10}$$

Then equation (9) reduces to:

$$\frac{\partial H_T}{\partial t} = D(S) \nabla^2 H_T - \gamma_H H_T \quad D(S) = D_H \left[ 1 + \left( \frac{D_{HS}}{D_H} - 1 \right) \frac{S}{K_D + S} \right] \tag{11}$$

Adding equations (6) and (7) yields an equation for total Scube:

$$\frac{\partial S_T}{\partial t} = D_S \nabla^2 S + D_{HS} \nabla^2 C + \alpha_S \frac{K_H^{n_H}}{K_H^{n_H} + (H_T)^{n_H}} - \gamma_S S \tag{12}$$

From assumption (3), changes in Scube concentration due to binding to Hedgehog to form a complex are negligible compared to diffusion and production of Scube:

$$\frac{\partial S_T}{\partial t} \approx D_S \nabla^2 S_T + \alpha_S \frac{K_H^{n_H}}{K_H^{n_H} + (H_T)^{n_H}} - \gamma_S S_T \tag{13}$$

Finally, the fast-equilibrium condition at the source leads to Hedgehog and Hedgehog-Scube complex fluxes:

$$\Phi_H = \kappa_H \frac{K_D}{S(0) + K_D} H_0 \quad \Phi_C = \kappa_H \frac{S(0)}{S(0) + K_D} H_0 \tag{14}$$

Hence the total Hedgehog flux is:

$$\Phi = \Phi_0 \left[ 1 + \left( \frac{\kappa_{HS}}{\kappa_H} - 1 \right) \frac{S(0)}{K_D + S(0)} \right] \tag{15}$$

Thus, under these assumptions, the total Hedgehog concentration obeys an ordinary diffusion-first-order-degradation equation with a *modified* diffusion constant and flux.

Although we have derived equations (11) and (15) under specific assumptions about the properties of Scube, the derivation motivates the general "phenomenological" model of the effect of Scube on Hedgehog in parameter regimes where the specific assumptions may not apply. In the following analysis, we will use these "phenomenological" equations to simplify our model.

### 2 Modeling setup

#### 2.1 Modeling framework

We model Hedgehog gradient formation in one dimension across a patterning field that initially has length 300 $\mu$m(Li et al., 2018). The boundary conditions are reflective at both ends for all species. The initial conditions are zero HH and Scube concentration throughout the field. At time $t = 0$, we turn on a scube-dependent flux $\Phi(S)$ at the left side of the patterning field. We then run all simulations until $t = 24$ hrs, by which time most models have reached steady-state.

In order to measure how well a particular model scales, we ran the model with a patterning field of length 200 $\mu$m, then plotted the gradient on the relative length $x/L$. For a model that scales perfectly, the gradient profiles should overlap exactly when plotted on $x/L$.

#### 2.2 Scaling metrics

To measure scaling, we consider two scaling metrics. To measure the shift in domain boundaries patterned by a gradient, we consider a previously described metric, which we refer to as "domain" metric (Ben-Zvi and Barkai, 2010). Three thresholds corresponding to three domain boundaries at positions $x/L = 0.25,\ 0.5,\ 0.75$ were determined from the gradient profile with $L = 300$ $\mu$m. We then computed the relative shift in each boundary $\delta_i$ when the field size was reduced to $L = 238$ $\mu$m. The "domain" metric was the average of these shifts:

$$\eta_{domain} = \frac{1}{3}\sum_{i=1}^{3}\delta_i \tag{16}$$

We also considered a scaling metric to measure scaling of the entire gradient profile. We computed the mean squared error between the gradient profiles in the long and short field sizes when plotted on the relative length $\bar{x} = x/L$ and normalized to the long gradient profile:

$$\eta_{profile} = \frac{\int_0^1 [H_{long}(\bar{x}) - H_{short}(\bar{x})]^2 d\bar{x}}{\int_0^1 H_{long}(\bar{x})^2 d\bar{x}} \tag{17}$$

This "profile" scaling metric is a more stringent condition than the "domain" metric, because profile metric requires that the gradient scales throughout the patterning field. In particular, for a model to scale with respect to the profile metric, the gradient amplitude must remain invariant to changes in patterning field size.

##### 2.3 Parameter gridsearch

We performed a parameter screen over various parameters in our model. Because the apparent diffusion constant of HH is relatively well characterized, we fix the diffusion constant with the value 1 $\mu$m$^2$/s (Li et al., 2018). We also fix the degradation rate $\gamma_H = 0.03$ min$^{-1}$ so that the lengthscale of the gradient matches approximately with the observed gradients in the zebrafish embryos. By fixing these two parameters, most of the gradients we consider are biologically reasonable.

We also define the scaled Hedgehog and Scube concentrations:

$$\bar{H} = \xi \frac{H}{\kappa H_0}, \quad \bar{S} = \frac{S}{K_D} \tag{18}$$

where $\xi = 1$ $\mu m$/min, and is defined to make the units of $\bar{H}$ dimensionless. Then the model equations reduce to the following form:

$$\frac{\partial \bar{H}}{\partial t} = D(S)\nabla^2 \bar{H} - \gamma_H \bar{H} \tag{19}$$

$$\frac{\partial \bar{S}}{\partial t} = D_S \nabla^2 \bar{S} + \alpha_S \frac{K_H^{n_H}}{K_H^{n_H} + \bar{H}^{n_H}} - \gamma_S \bar{S} \tag{20}$$

$$D(S) = D_H \left[ 1 + \left( \frac{D_{HS}}{D_H} - 1 \right) \frac{\bar{S}}{1 + \bar{S}} \right] \tag{21}$$

$$D(S) \cdot \left. \frac{\partial \bar{H}}{\partial x} \right|_{x=0} = 1 + \left( \frac{\kappa_{HS}}{\kappa_H} - 1 \right) \frac{\bar{S}(0)}{1 + \bar{S}(0)} \tag{22}$$

We conducted a parameter screen on all remaining parameters in the model, varying each parameter systematically over several orders of magnitude. Table 2 gives the parameter grid used in the gridsearch. For each combination of the $\sim 40,000$ parameters we scanned, we ran the models for the two lengths of the patterning field and computed both the profile and domain scaling metrics between the ligand distribution $H(x)$.

### 3 Uniform first-order degradation of Hedgehog

Panels A-G from supplementary figure 7 show the distribution of scaling metrics for fixed values of each parameter scanned in the model (Fig. S7). The panels in the figure show the distribution of the domain metric, and the distribution of the profile metric was qualitatively very similar. The results display many features that are reminiscent of the classic expansion-repression model (Fig. S7A-E): scaling works best when Scube diffusion is fast, Scube degradation is not too fast, and Scube repression by Hedgehog is sharp (Ben-Zvi and Barkai, 2010). Additionally, scaling works best when the threshold for Scube repression

| Parameter Names | Symbols | Units |
|---|---|---|
| Hedgehog diffusion constant | $D_H$ | $\mu m^2$/min |
| Hedgehog degradation rate | $\gamma_H$ | $\text{min}^{-1}$ |
| Scaled Scube-Hedgehog repression threshold | $K_H$ | dimensionless |
| Scaled Scube production rate | $\alpha_S$ | $\text{min}^{-1}$ |
| Scube degradation rate | $\gamma_S$ | $\text{min}^{-1}$ |
| Scube-Hedgehog repression hill coefficient | $n_H$ | dimensionless |
| Scube diffusion constant | $D_S$ | $\mu m^2$/min |
| Relative increase in diffusion constant | $D_{HS}/D_H$ | dimensionless |
| Relative increase in release rate | $\kappa_{HS}/\kappa_H$ | dimensionless |
| Relative increase in degradation rate | $\gamma_{HS}/\gamma_H$ | dimensionless |
| Second order Hedgehog degradation rate | $\gamma_{H2}$ | $\mu\text{M}^{-1} \cdot \text{min}^{-1}$ |
| Hedgehog flux | $\Phi_0$ | $\mu\text{M} \cdot \text{min}^{-1} \cdot \mu m^{-2}$ |

Table 1: Parameter names and symbols

| | |
|---|---|
| $K_H$ | $\{10^{-5}, 3 \cdot 10^{-5}, 10^{-4}, 3 \cdot 10^{-4}, 10^{-3}, 3 \cdot 10^{-3}, 10^{-2}, 3 \cdot 10^{-2}, 10^{-1}\}$ |
| $\alpha_S$ | $\{10^{-4}, 3 \cdot 10^{-4}, 10^{-3}, 3 \cdot 10^{-3}, 10^{-2}, 3 \cdot 10^{-2}, 10^{-1}, 3 \cdot 10^{-1}, 1\}$ |
| $\gamma_S$ | $\{0, 10^{-3}, 10^{-2}, 10^{-1}\}$ |
| $n_H$ | $\{1, 3, 10\}$ |
| $D_S$ | $\{10, 100, 1000\}$ |
| $D_{HS}/D_H$ | $\{1, 3, 10, 30\}$ |
| $\kappa_{HS}/\kappa_H$ | $\{1, 3, 10, 30\}$ |

Table 2: Ranges for parameters in uniform-degradation parameter gridsearch

by Hedgehog is not to low. Strikingly, we see that in the uniform-degradation case, both an increase in the diffusion constant *and* an increase in the release rate is required for scaling (Fig. S7F-G).

Next, we fixed the parameters $K_H$, $\alpha_S$, $\gamma_S$, $n_H$, and $D_S$ to produce a model that could scale under certain conditions, and generated a "phase-plot" to ask how the scaling depended quantitatively on the two remaining free parameters: the relative increase in release rate $\kappa_{HS}/\kappa_H$ and diffusion constant $D_{HS}/D_H$. We found that models that scale best balance an increase in release rate with an increase in the diffusion constant.

To explain this, consider a simplified model of gradient formation. Suppose Hedgehog diffuses through the field with diffusion constant $D$ and is removed at rate $\gamma$. Given a constant flux boundary condition at the left side of the patterning field and the condition that the right side of the patterning field is much greater than the lengthscale $\lambda$, the steady state Hedgehog concentration is:

$$H(x) = \frac{\Phi}{\sqrt{\gamma D}} \exp(-x/\lambda), \quad \lambda = \sqrt{\frac{D}{\gamma}} \tag{23}$$

In order for a gradient to perfectly scale, the amplitude must remain invariant and the lengthscale must increase. Scube cannot increase the lengthscale of the gradient by increasing the flux and hence must increase the diffusion constant. However, by increasing the diffusion constant, the amplitude at the origin decreases. In order to keep the amplitude invariant, Scube must also increase the flux to compensate. Hence, both actions are required for scaling in this simple model.

### 4 Self-enhanced degradation of Hedgehog

In the Hedgehog system, one major route of Hedgehog degradation is endocytosis mediated by the PTCH receptor. Hedgehog signaling upregulates the production of PTCH and hence increases HH degradation(Li et al., 2018). We consider a simplified model that captures the fact that Hedgehog signalling increases the degradation rate of HH. We replace the degradation term with a second-order degradation term $\gamma_H H^2$. Because of the second-order degradation term, we must use the equation for the absolute Hedgehog concentration rather than the scaled Hedgehog equation. The equation for Hedgehog is:

$$\frac{\partial H}{\partial t} = D(S)\nabla^2 H - \gamma_{H2} H^2 \quad D(S) = D_H \left[1 + \left(\frac{D_{HS}}{D_H} - 1\right) \frac{S}{K_D + S}\right] \tag{24}$$

Self-enhanced degradation has been implicated in scaling (Ben-Zvi and Barkai, 2010; Eldar et al., 2003; Lander et al., 2009). This is because a self-enhanced degradation scheme confers robustness to changes in morphogen flux rate to the gradient in the region distal to the morphogen source.

In order to see this, consider a simplified system without Scube. At steady state, the morphogen distribution $H(x)$ satisfies:

$$\frac{d^2H}{dx^2} = \gamma_{H2}H^2$$

The solution to this equation subject to the condition that $H(x) \to 0$ as $x \to \infty$ is:

$$H(x) = \frac{6D}{\gamma_{H2}(x+\epsilon)^2}$$

Requiring that $DH' = -\Phi$ yields:

$$\epsilon = \sqrt[3]{\frac{12D^2}{\Phi\gamma_{H2}}}$$

If $\epsilon$ is small in comparison with the size of the patterning field, then $H(x) \approx 6D/\gamma_{H2}x^2$. Note that in this regime, the morphogen distribution does not depend on the flux into the system. In particular, the gradient can now be scaled for $x \gg \epsilon$ by increasing the diffusion constant only. Scaling will occur in the region far from the morphogen source $x \gg \epsilon$, but scaling at the side of the patterning field near the producing cells does not occur if scube modulates the diffusion constant only.

We ran the model with the self-enhanced degradation term over the same parameter gridsearch as the uniform degradation case. We consider two parameter regimes: a "high production" regime where $\Phi$ is large enough so that $\epsilon$ is small in comparison to the length of the patterning field, and a "low production" regime where $\Phi$ is small enough so that $\epsilon$ is comparable to the length of the patterning field.

In the "low production" regime, the results are qualitatively similar to the uniform degradation case. Because the self-enhanced degradation does not provide the system with robustness to changes in the morphogen flux, both the release and expansion interactions were required to produce scaling (Fig. S8A-D).

In contrast, in the "high production" regime, only the expansion interaction was required to produce scaling under the domain scaling metric (Fig. S8, panels E-H). This is because the domain scaling metric only requires the gradient to scale at particular points within the patterning field: $L/4$, $L/2$, and $3L/4$. If $L \gg \epsilon$, then the gradient scales at these points. In particular, the domain scaling metric does not require that the gradient amplitude remains invariant as the patterning field size is changed. Indeed, under the profile metric that does require that the amplitude remain invariant, scaling requires both the release and expansion interactions (Fig. S8I-J).

Thus, even though in the self-enhanced degradation scheme, under certain conditions the release interaction may not be strictly required, the release interaction serves at least two purposes. First, it allows the system to scale in a wider parameter regime - models with the release interaction continue to scale even when the morphogen production rate is decreased to a point where the robustness conferred by self-enhanced degradation is lost (Fig. 7G in the main text). Second, it allows the system to fix the morphogen gradient amplitude when the patterning field size is changed.

### 5 Heterozygous Sonic Hedgehog knockout

Knocking out one allele of Sonic Hedgehog leads to mice that develop normally (Chiang et al., 1996). It is unclear whether gene dosage affects the rate at which Hedgehog is secreted from cells. Nevertheless, because Scube2 can promote release of Hedgehog from Hedgehog-secreting cells, we wondered if Scube2-mediated feedback could be a mechanism to explain the robustness of tissue patterning to changes in the Hedgehog gene dosage. To investigate this question, we one again conducted a parameter screen using the uniform-degradation model discussed in section 3 on the set of $\sim 40,000$ parameters in Table 2. To quantify whether a particular model was robust to changes in Hedgehog flux, we ran each model twice, halving the Hedgehog flux on the second run. Similar to the profile metric, we used the ligand distribution $H(x)$ to draw three domain boundaries, then computed the average shift in domain boundary. More robust models have smaller values for this metric.

The models that lead to flux-robustness generally have a large value for $\kappa_{HS}/\kappa H$, and a small value for $D_{HS}/D_H$ (Fig. S9). This suggests that the release interaction is important for flux-robustness, but the diffusion interaction may in fact hinder flux-robustness. The release interaction is important in fixing the gradient amplitude. In this case, because the flux of Hedgehog is halved without increasing the patterning field length, the diffusion interaction may actually interfere with flux-robustness by increasing the lengthscale of the gradient when it does not need to be increased.

Overall, these results suggest that although it is unlikely that Scube feedback alone can explain the robustness of tissue patterning to changes in HH gene dosage, Scube feedback can in certain regimes confer flux robustness to the HH gradient in addition to scaling it.

### 6 Scube modulating Hedgehog degradation

In the classic expansion repression model, the expander can expand the morphogen gradient in two ways: it can increase the diffusion constant of the morphogen or decrease the degradation rate of the morphogen. Up until this point, based upon the mechanism by which the Scube2 protein is thought to act, we have considered models where Scube2 modulates the diffusive and release properties of Hedgehog only.

Here we consider models where Scube2 modulates the *degradation* and release properties of Hedgehog. We begin with a brief theoretical motivation. As discussed previously, the steady-state Hedgehog distribution in the case of uniform degradation is:

$$H(x) = \frac{\Phi}{\sqrt{\gamma D}} \exp(-x/\lambda),\ \lambda = \sqrt{\frac{D}{\gamma}}$$

In order to expand the range of the morphogen, Scube2 must decrease the degradation rate $\gamma$. If Scube2 expands the Hedgehog gradient by decreasing the HH degradation rate, in order to leave the amplitude invariant, Scube2 must also decrease the release rate from the HH secreting cells, which is inconsistent with observed biochemical data (Tukachinsky et al., 2012; Creanga et al., 2012).

We ran the same gridsearch as previously described, allowing Scube2 to modulate the release rate or the *degradation* rate of Hedgehog. Models that led to the best scaling involve Scube2 decreasing both the degradation rate and the HH flux (Fig. S10). Because biochemical evidence suggests that Scube2 *promotes* HH release rather than inhibiting it, our results suggest that Scube2 is unlikely to expand the HH gradient by decreasing degradation rate of HH.

### 7 Hedgehog flux scales with patterning field size

We now consider a proposed model for scaling by scaling the morphogen source size. When the size of an animal is decreased, it is possible that the number of ligand-producing cells is correspondingly decreased. It is unclear whether a decrease in the number of HH-producing cells will actually lead to a decreased flux of Hedgehog into the patterning field. In particular, if the rate limiting step in Hedgehog flux is the release of Hedgehog off the membranes of ligand-producing cells, increasing the number of cells may not necessarily increase the ligand flux into the system. Nevertheless, we considered models where the flux of Hedgehog changes with the size of the patterning field.

We consider models where the HH flux into the field in the absence of Scube is proportional to the size of the patterning field (Fig. S11). Once again, we ran a parameter gridsearch to determine which models led to scaling. Our results show that the scaling of the morphogen source size cannot explain the observed scaling behavior alone. Because the lengthscale of the gradient depends on the diffusion constant and not the rate of Hedgehog flux, the gradient cannot be scaled by changing the Hedgehog flux only. However, notably the release interaction is no longer required for scaling to occur, because the amplitude problem is fixed by the increased flux as the patterning field size increases.

**Supplemental Figures:**

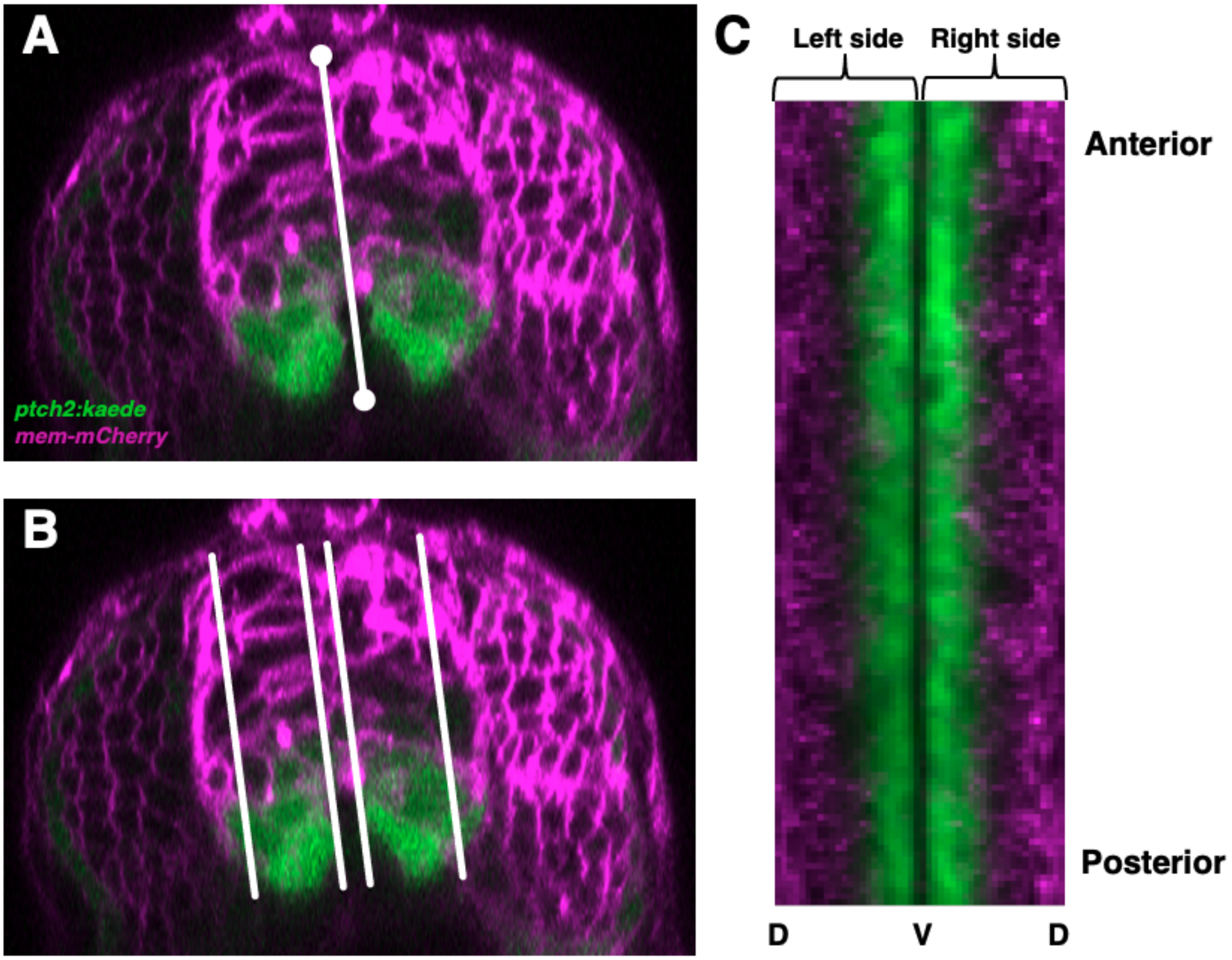


**Figure S1- Segmentation of neural imaging data.**

(A) Image of a 20 hpf *Tg(ptch2:kaede)* reporter embryo undergoing selection of the “axis of reflection” which serves to mark a measurement of Dorsoventral height and separate the left and right halves of the neural tube. These positions are picked by the user, first by picking the bottom of the floor plate cell, then inputting the top coordinate of the roof plate cell. (B) Image of a 20 hpf *Tg(ptch2:kaede)* reporter embryo after a user has selected the width of the spinal cord. The algorithm then calculates how much imaging data to collect based on a ratio which avoids mature neurons and the lumen of the spinal cord. (C) After collection of average intensities in each bin, data is stored as shown. Average profiles for generating distribution plots and segmenting domains are gathered by averaging these data along the A-P axis for both halves of the neural tube.

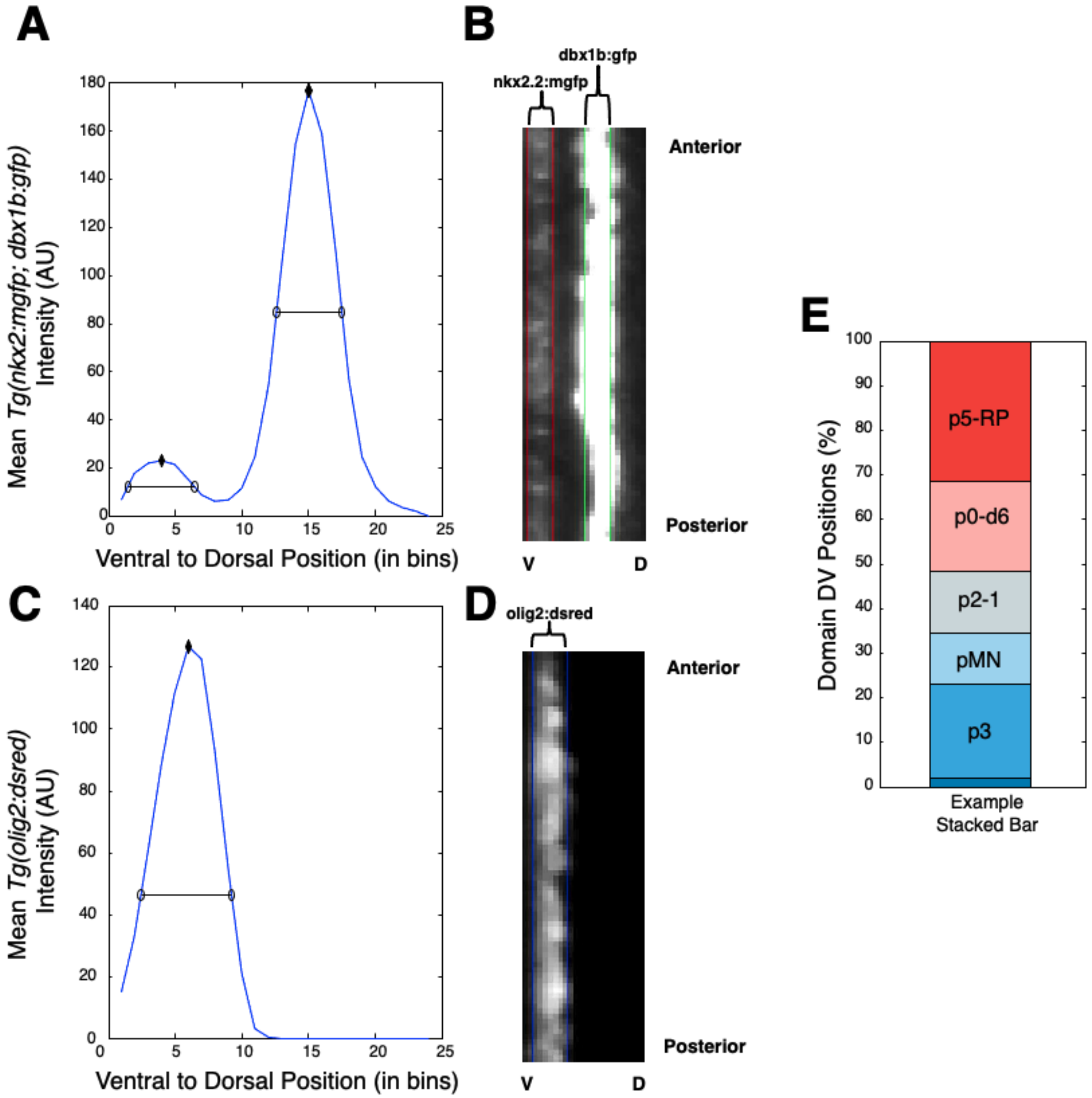


**Figure S2- Progenitor domain width determination and bar plot generation.**

(A) Averaged image intensity profiles from the green channel of both sides of a segmented neural tube from a *Tg(dbx1b:GFP;olig2:dsred;nkx2.2a:memGFP)* embryo. Black diamonds represent peaks found by a peak finding algorithm, while open circles and lines show the calculated domain boundaries and width for this embryo based on the universally applied threshold in this dataset. Thresholds are determined by 50% of average peak intensity of the control population for each dataset. (B) Example domain determination of *nkx2.2a* and *dbx1b*+ cells. Red lines mark the predicted *nkx2.2a* domain, which correlates with the boundary of their fluorescence. Green lines mark the predicted width of the *dbx1b* domain which correlates well with visible fluorescence of this domain. Some anterior-posterior variability in domain size is observed. (C) Formatted as in part A, this plots the averages *olig2:dsred*+ intensity, peak, and determined width. (D) Example domain determination of *olig2*+ cells. Blue lines mark the predicted *olig2:dsred* domain, which correlates with the boundary of their fluorescence. Thresholds are determined by 25% of average peak intensity of the control embryos in each dataset. (E) Example stacked bar plot generated only from this embryo using calculated domain positions to determine domain sizes.

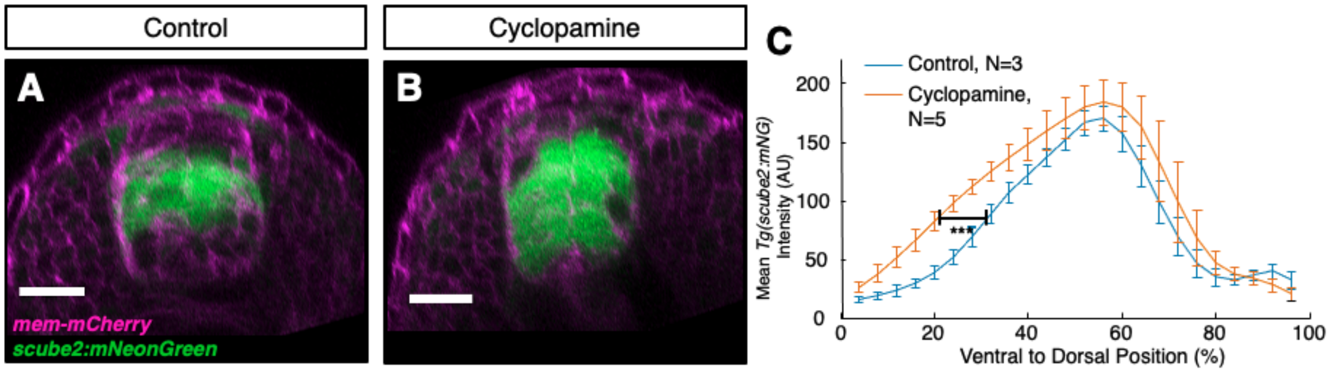


**Figure S3- Cyclopamine treatment of *tg(scube2:moxNG)* embryos.**

(A-C) Transverse view 22 hpf *Tg(scube2:moxNG*; *mem:mCherry)* embryos treated with a DMSO control (A) or 100uM Cyclopamine (B). (C) Quantification of mean reporter intensity of embryos as treated in A-B. The black bar marks the position of 50% of control maximum intensity which was used for statistical testing. These values were statistically significantly shifted ventrally in drug treated embryos relative to control (unpaired t-test p= P=.0001145.).

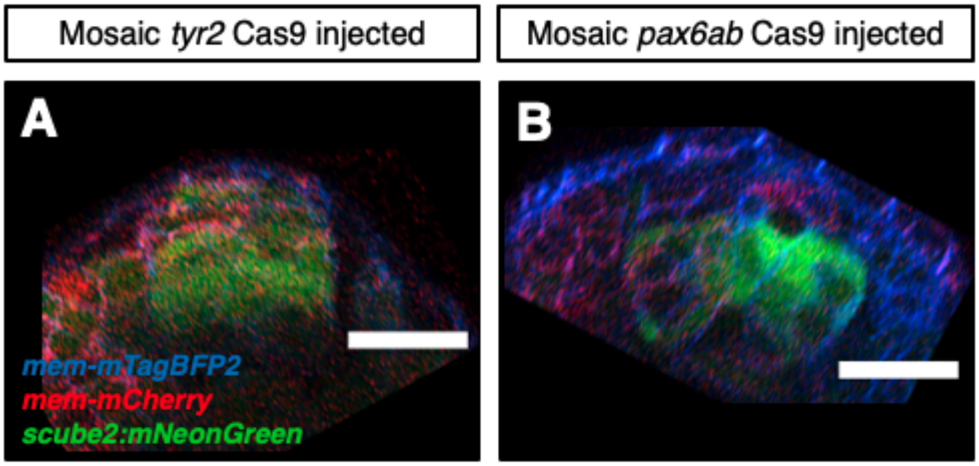


**Figure S4- Mosaic *pax6a/b* CRISPR mutants have lowered *scube2* expression.**

(A-B) *Tg(scube2:moxNG)* embryos imaged at 18 hpf that were injected at the single cell stage with *mem-mTagBFP2* mRNA and injected at the 8-16 cell stage with *mem-mCherry* mRNA, Cas9 protein, and sgRNAs targeting either the tyrosinase pigment gene as a control (A) or *pax6a* and *b* (B).

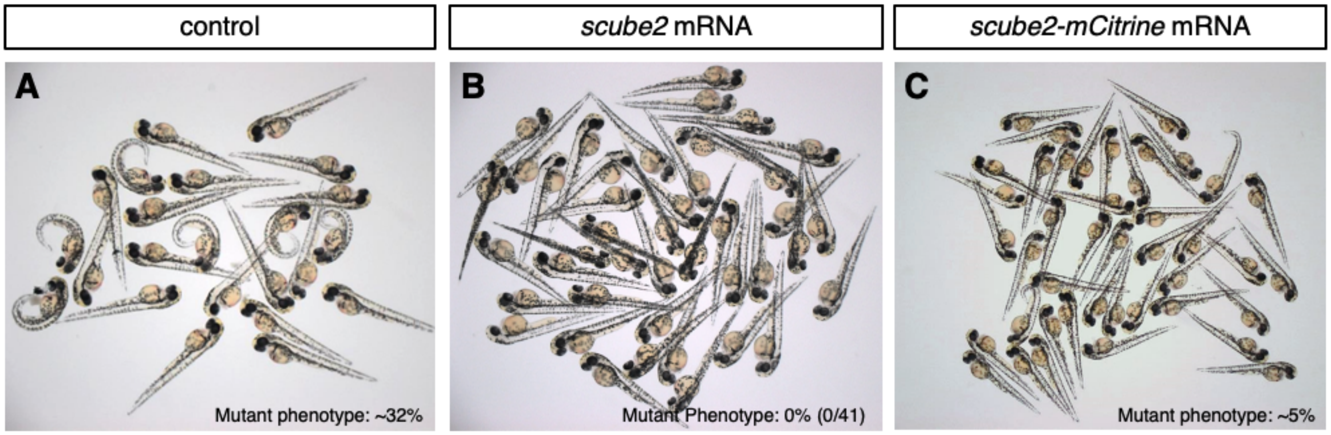


**Figure S5- Rescue of scube2 CRISPR mutants with *scube2* or *scube2-mCitrine* mRNA.**

(A) Results of a *scube2* mutant in-cross. The allele was generated by mutagenesis with CRISPR using three guides targeting *scube2* coding sequence (B) Embryos rescued by the injection of *scube2* mRNA co-injected with *mem-mCardinal* which were screened for being mem-mCardinal positive (Liu et al., 2018). (C) Embryos rescued by the injection of *scube2-mCitrine* mRNA co-injected with *mem-mCardinal* which were also screened for *mem-mCardinal* fluorescence.

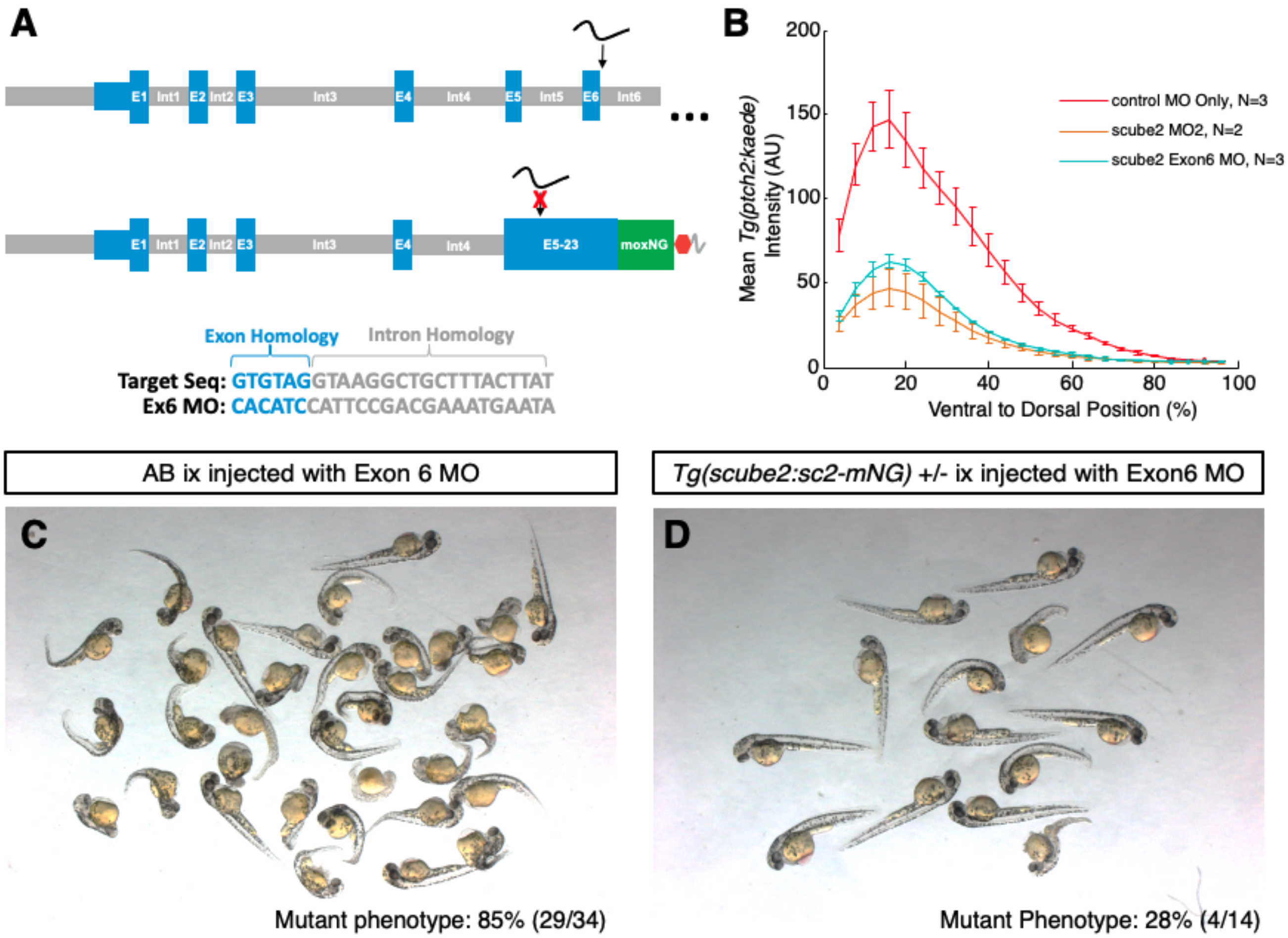


**Figure S6- Rescue of morpholino knockdown by *tg(scube2:scube2-moxNeonGreen)* expression.**

(A) Schematic of Scube2 Exon 6 targeting morpholino and expected resistance in the genome of *tg(scube2:scube2-moxNeonGreen)* embryos. Splice junction targeted by the Exon 6 morpholino is largely absent from the transgenic full length Scube2-mCitrine construct, allowing a test of rescue capacity. (B) *tg(ptch2:kaede)* response profiles of embryos injected wither with p53 morpholino only, the previously published Scube2 MO2 (Woods and Talbot, 2005), or Scube2 Exon 6 morpholino. (C) Overview photo of wildtype embryos injected with Scube2 Exon6 morpholino showing the tail curling indicative of Shh signaling phenotypes. (D)

Overview photo of *tg(scube2:scube2-moxNeonGreen)* embryos injected with Scube2 Exon6 morpholino.

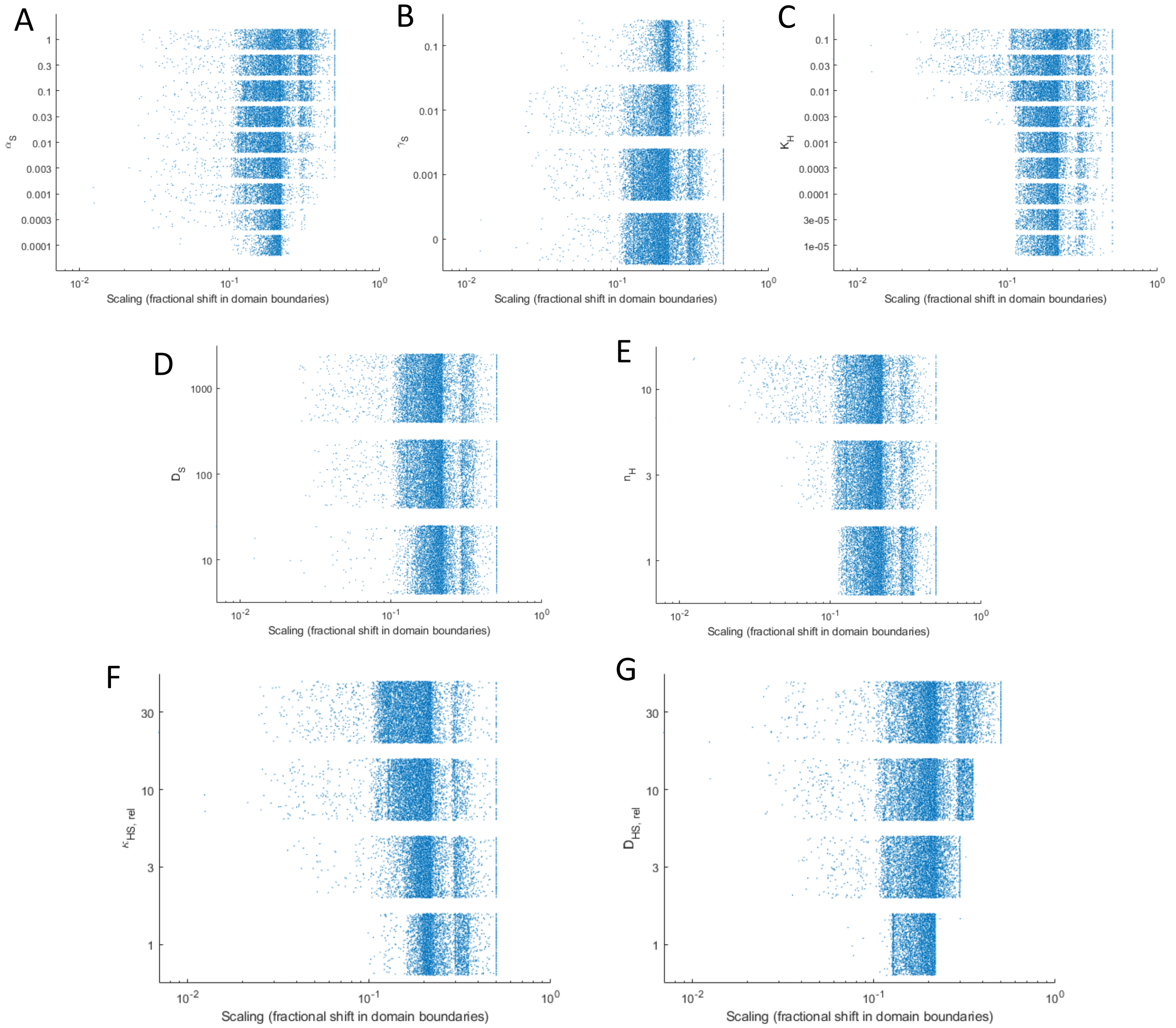


H

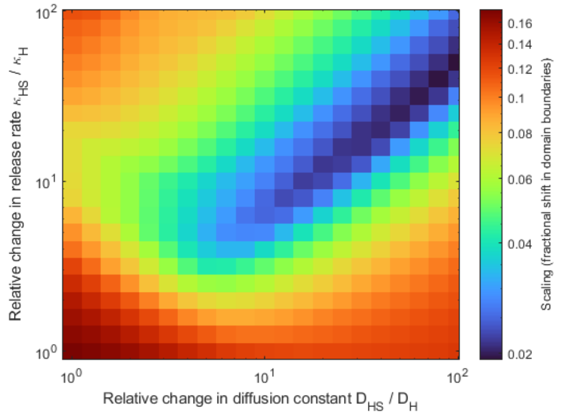


I

| | |
|---|---|
| $K_H$ | 0.03 |
| $\alpha_S$ | 0.01 $\text{min}^{-1}$ |
| $\gamma_S$ | 0.003 $\text{min}^{-1}$ |
| $n_H$ | 10 |
| $D_S$ | 1000 $\mu m^2/\text{min}$ |

**Figure S7- Uniform degradation model.**

(A-G) Values of scaling metric (as measured by shift in domain boundaries) produced by a set of parameters. Each blue dot is a simulation. For each panel, a different parameter is changed. (A) Scaled Sc2 production rate, αS. (B) Sc2 degradation rate, γS. (C) Scaled Sc2-Hh repression threshold, KH. (D) Sc2 diffusion constant, DS. (E) Sc2-Hh repression hill coefficient, nH. (F) relative increase in release rate, κHS/κH. (G) Relative increase in diffusion constant, DHS/DH. (H)

Phase plot of scaling measured by fractional shift in domain boundaries across relative changes in release rate and in diffusion constant. (I) Parameters that were used in the model to produce phase plot in (H).

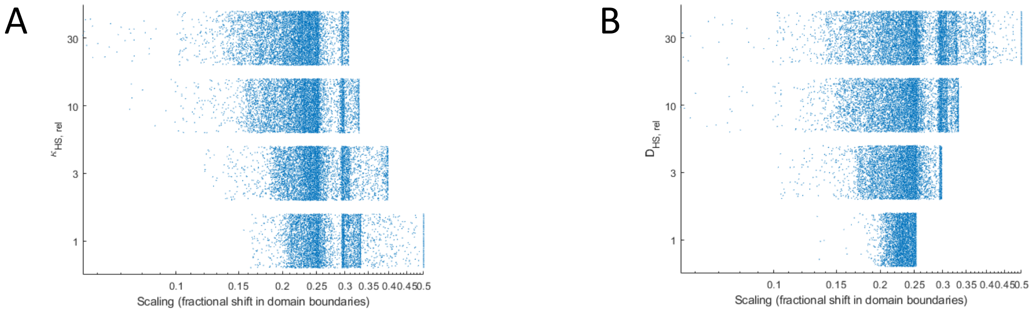


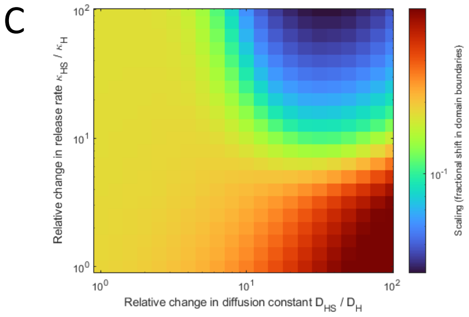


D

| | |
|---|---|
| $K_H$ | $0.03\ \mu$M |
| $\alpha_S$ | $0.1\ \text{min}^{-1}$ |
| $\gamma_S$ | $0.001\ \text{min}^{-1}$ |
| $n_H$ | 10 |
| $D_S$ | $1000\ \mu m^2/\text{min}$ |
| $\gamma_{H2}$ | $1\ \mu\text{M}/\text{min}$ |
| $\Phi_0$ | $0.1\ \mu\text{M}/\text{min} \cdot \mu m^2$ |

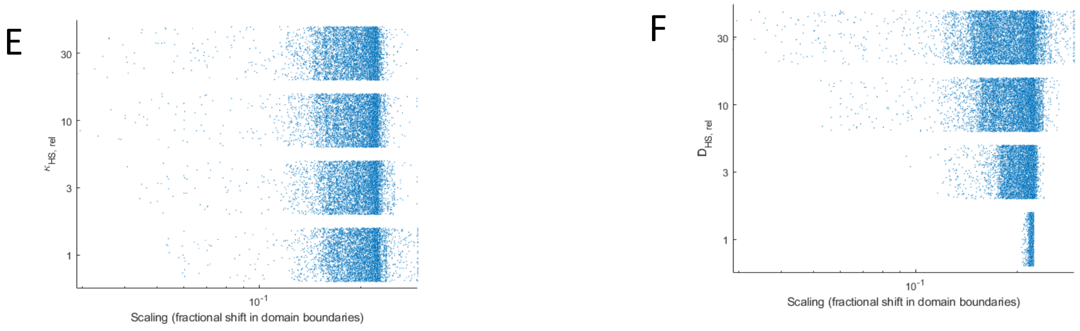


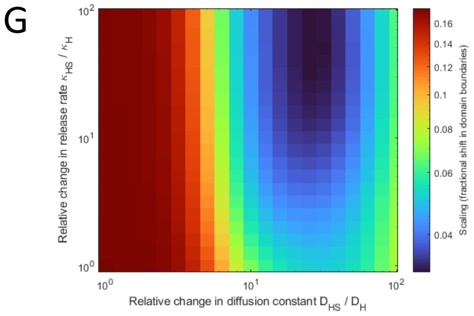


H

| | |
|---|---|
| $K_H$ | $0.03\ \mu$M |
| $\alpha_S$ | $0.1\ \text{min}^{-1}$ |
| $\gamma_S$ | $0.001\ \text{min}^{-1}$ |
| $n_H$ | 10 |
| $D_S$ | $1000\ \mu m^2/\text{min}$ |
| $\gamma_{H2}$ | $1\ \mu\text{M}/\text{min}$ |
| $\Phi_0$ | $30\ \mu\text{M}/\text{min} \cdot \mu m^2$ |

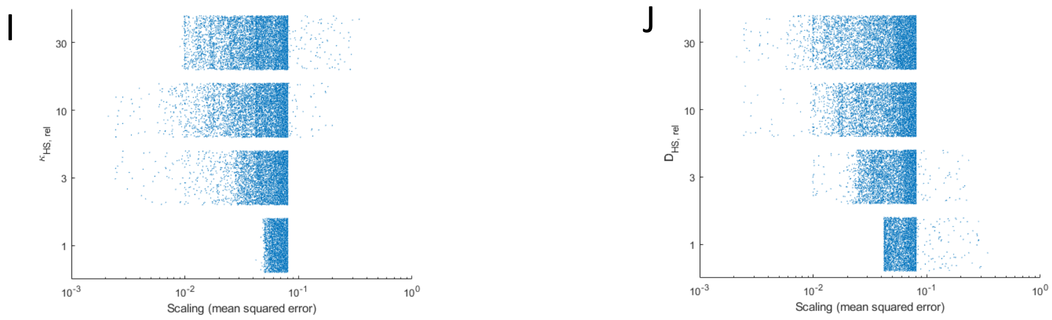

**Figure S8 - Self-enhanced degradation model.**

(A-F) Plots from self-enhanced degradation model where there is low production of hedgehog. (A, B, E, F) Values of scaling metric (as measured by shift in domain boundaries) produced by a set of parameters. For each panel, a different parameter is changed. (A) Each blue dot is a simulation generated by different values of one parameter, relative increase in release rate, κHS/κH (Y-axis), while other parameters are kept constant, that produce certain scaling metric values (X-axis). (B) Each blue dot is a simulation generated by different values of one parameter, relative increase in diffusion constant, DHS/DH (Y-axis), while other parameters are kept constant, that produce certain scaling metric values (X-axis). (C) Phase plot of scaling measured by fractional shift in domain boundaries across relative changes in release rate and in diffusion constant. (D) Parameters that were used in the model to produce phase plot in (C). (E) Each blue dot is a simulation generated by different values of one parameter, relative increase in diffusion constant, relative increase in release rate, κHS/κH (Y-axis), while other parameters are kept constant, that produce certain scaling metric values (X-axis). (F) Each blue dot is a simulation generated by different values of one parameter, relative increase in diffusion constant, relative increase in diffusion constant, DHS/DH (Y-axis), while other parameters are kept constant, that produce certain scaling metric values (X-axis). (G-J) Plots from self-enhanced degradation model where there is high production of hedgehog. (G) Phase plot of scaling measured by fractional shift in domain boundaries across relative changes in release rate and in diffusion constant. (H) Parameters that were used in the model to produce phase plot in (G). (I) Each blue dot is a simulation generated by different values of one parameter, relative increase in release rate, κHS/κH (Y-axis), while other parameters are kept constant, that produce certain scaling metric values (X-axis). (J) Each blue dot is a simulation generated by different values of one parameter, relative increase in diffusion constant, relative increase in diffusion constant, DHS/DH (Y-axis), while other parameters are kept constant, that produce certain scaling metric values (X-axis).

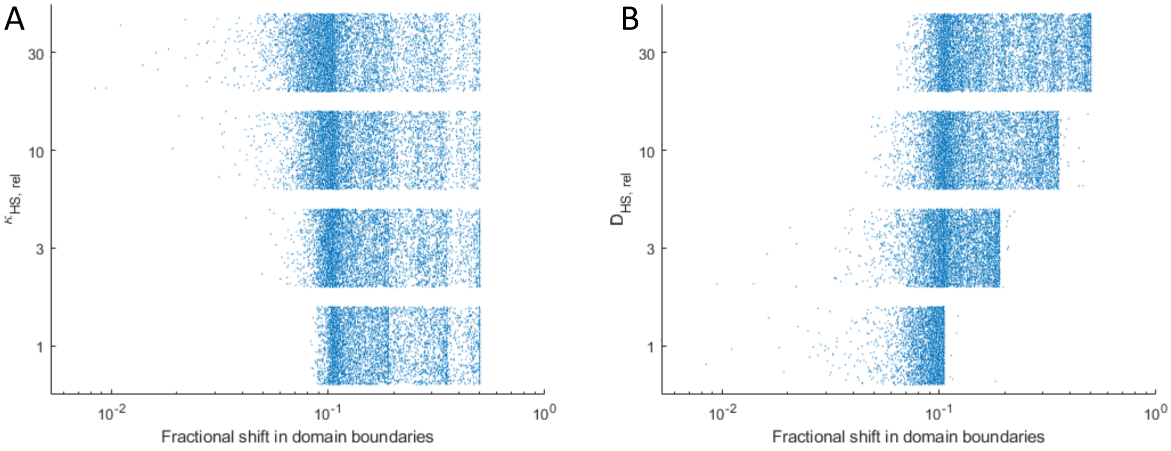


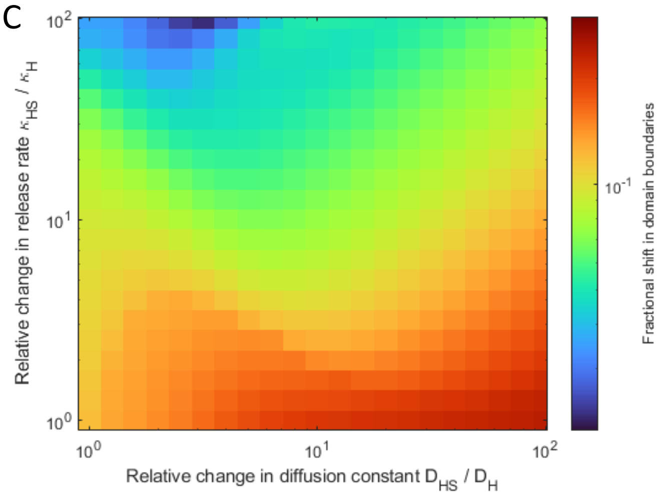


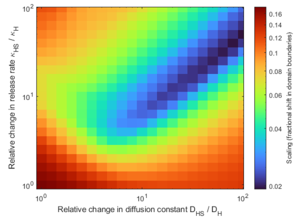

| | |
|---|---|
| $K_H$ | 0.03 |
| $\alpha_S$ | 0.01 $\mathrm{min}^{-1}$ |
| $\gamma_S$ | 0.003 $\mathrm{min}^{-1}$ |
| $n_H$ | 10 |
| $D_S$ | 1000 $\mu m^2/\mathrm{min}$ |

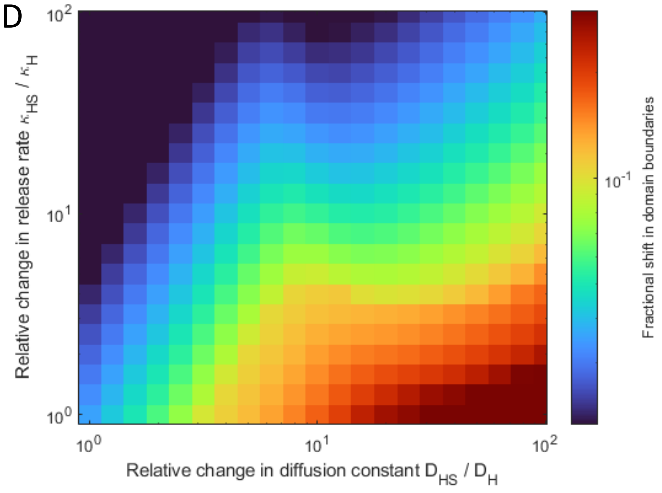


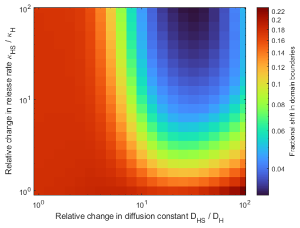

| | |
|---|---|
| $K_H$ | 0.03 $\mu$M |
| $\alpha_S$ | 0.1 $\mathrm{min}^{-1}$ |
| $\gamma_S$ | 0.001 $\mathrm{min}^{-1}$ |
| $n_H$ | 10 |
| $D_S$ | 1000 $\mu m^2/\mathrm{min}$ |
| $\gamma_{H2}$ | 1 $\mu$M/min |
| $\Phi_0$ | 1 $\mu\mathrm{M}/\mathrm{min}\cdot\mu m^2$ |

**Figure S9- Effect of 50% reduction in Hedgehog flux in uniform degradation and self-enhanced degradation models.**

(A) Each blue dot is a simulation generated by different values of one parameter, relative increase in diffusion constant, DHS/DH (Y-axis), while other parameters are kept constant, that produce certain scaling metric values (X-axis). (B) Each blue dot is a simulation generated by different values of one parameter, relative increase in release rate, κHS/κH (Y-axis), while other parameters are kept constant, that produce certain scaling metric values (X-axis). (C) Phase plot, for uniform degradation model, of scaling measured by fractional shift in domain boundaries across relative changes in release rate and in diffusion constant. (D) Smaller plot is the same as the phase plot in Figure 1. (E) Table shows the parameters that were used in the model to produce the phase plot. (F) Phase plot, for self-enhanced degradation model, of scaling measured by fractional shift in domain boundaries across relative changes in release rate and in diffusion constant. (G) Smaller plot is the same as the phase plot in Figure 2. (H) Table shows the parameters that were used in the model to produce the phase plot.

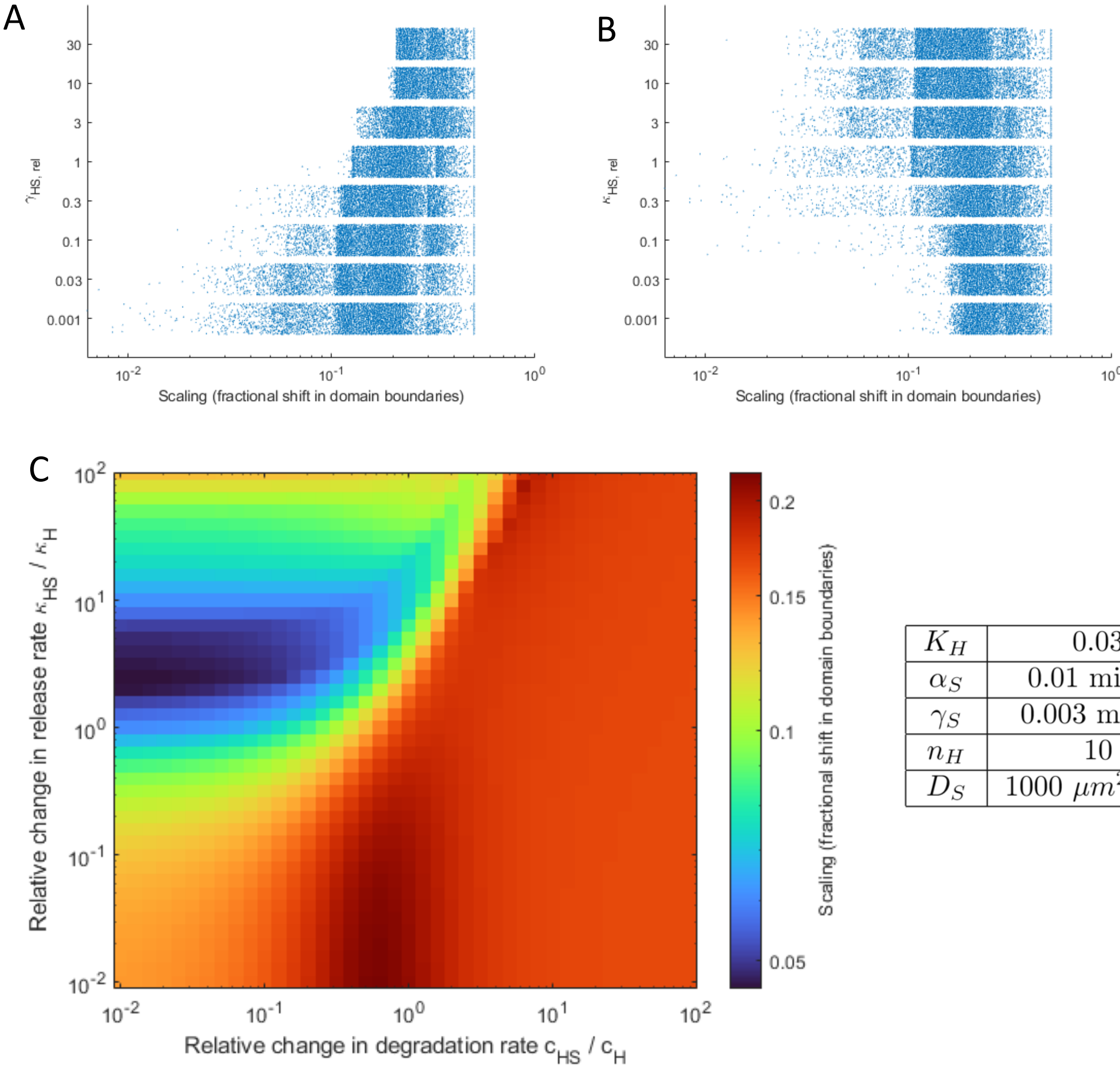


| | |
|---|---|
| $K_H$ | 0.03 |
| $\alpha_S$ | 0.01 $\mathrm{min}^{-1}$ |
| $\gamma_S$ | 0.003 $\mathrm{min}^{-1}$ |
| $n_H$ | 10 |
| $D_S$ | 1000 $\mu m^2/\mathrm{min}$ |

**Figure S10- Model in which Scube2 enhances degradation of Hedgehog.**

(A) Each blue dot is a simulation generated by different values of one parameter, relative increase in degradation rate, γHS/γH, while other parameters are kept constant, that produce certain scaling metric values (X-axis). (B) Each blue dot is a simulation generated by different values of one parameter, relative increase in release rate, κHS/κH (Y-axis), while other parameters are kept constant, that produce certain scaling metric values (X-axis). (C) Phase plot of scaling measured by fractional shift in domain boundaries across relative changes in release rate and in degradation rate. (D) Parameters that were used in the model to produce phase plot in (C).

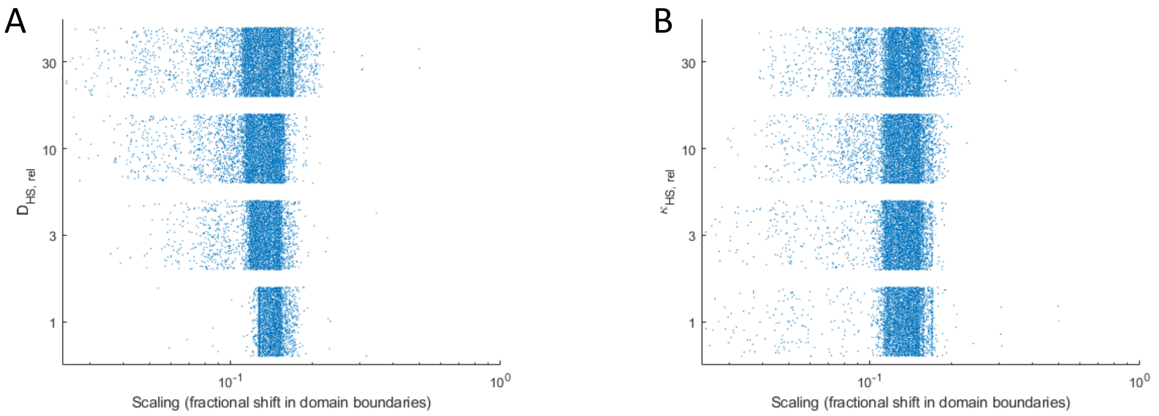


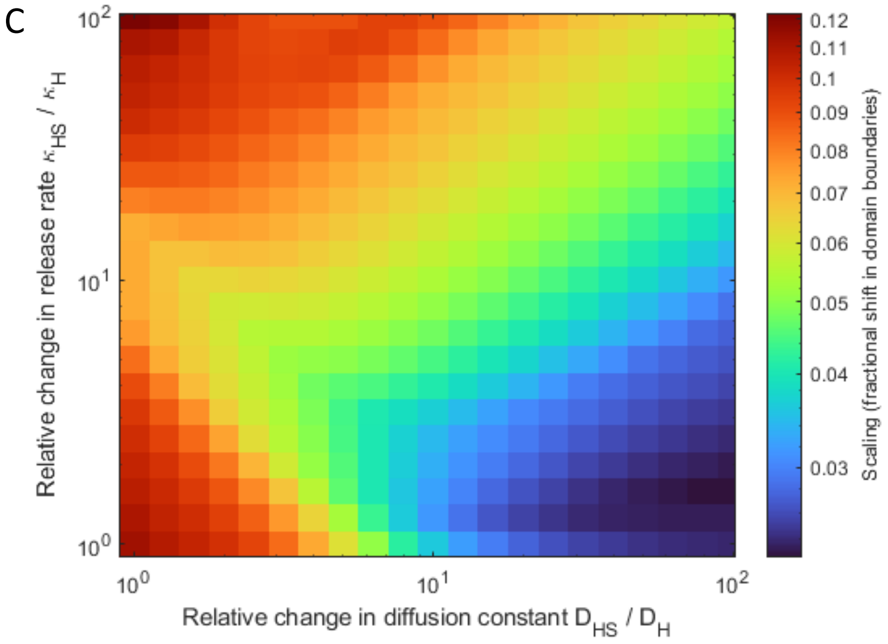


| | |
|---|---|
| $K_H$ | 0.03 |
| $\alpha_S$ | 0.01 $\text{min}^{-1}$ |
| $\gamma_S$ | 0.003 $\text{min}^{-1}$ |
| $n_H$ | 10 |
| $D_S$ | 1000 $\mu m^2/\text{min}$ |

**Figure S11- Model in which Hedgehog flux scales with the size of the patterning field.**

(A) Each blue dot is a simulation generated by different values of one parameter, relative increase in diffusion constant, DHS/DH (Y-axis), while other parameters are kept constant, that produce certain scaling metric values (X-axis). (B) Each blue dot is a simulation generated by different values of one parameter, relative increase in release rate, κHS/κH (Y-axis), while other parameters are kept constant, that produce certain scaling metric values (X-axis). (C) Phase plot of scaling measured by fractional shift in domain boundaries across relative changes in release rate and in diffusion constant. (D) Parameters that were used in the model to produce phase plot in (C)

**Supplemental Movies:**

**Movie S1-** ***Tg(scube2:moxNG; shha:mem-mCherry)*** **timelapse transverse view**

**Movie S2-** ***Tg(scube2:moxNG; shha:mem-mCherry)*** **timelapse maximum intensity projection dorsal view**

**Movie S3- Scube2-mCitrine FRAP imaging in a dome stage embryo**

**Movie S4- Sec-mCitrine FRAP imaging in a dome stage embryo**

**Movie S5-** ***Tg(scube2:scube2-moxNG; shha:mem-mCherry)*** **timelapse transverse view**

**Movie S6-** ***Tg(scube2:scube2-moxNG; shha:mem-mCherry)*** **timelapse maximum intensity projection dorsal view**
